# Persistent DNA damage promotes microglial dysfunction in Ataxia-telangiectasia

**DOI:** 10.1101/2021.06.02.446712

**Authors:** Julie Bourseguin, Wen Cheng, Emily Talbot, Liana Hardy, Svetlana V. Khoronenkova

## Abstract

The autosomal recessive genome instability disorder Ataxia-telangiectasia, caused by mutations in ATM kinase, is characterised by the progressive loss of cerebellar neurons. We find that DNA damage associated with ATM loss results in dysfunctional behaviour of human microglia, immune cells of the central nervous system. Microglial dysfunction is mediated by the pro-inflammatory RELB/p52 non-canonical NF-κB transcriptional pathway and leads to excessive phagocytic clearance of neurites. Pathological phagocytosis of neuronal processes by microglia has also been observed in multiple sclerosis, Alzheimer’s and progranulin deficiency, suggesting a common mechanism that promotes neuronal damage. Activation of the RELB/p52 pathway in ATM-deficient microglia is driven by persistent DNA damage and is dependent on the NIK kinase. These results provide insights into the underlying mechanisms of aberrant microglial behaviour in Ataxia-telangiectasia, potentially contributing to neurodegeneration.

## INTRODUCTION

Ataxia-telangiectasia (A-T) is an autosomal recessive disease caused by loss-of-function mutations in the *ATM* gene (A-T mutated) (1). ATM is a serine/threonine protein kinase that plays a central role in coordinating the cellular response to genotoxic stress, in particular cytotoxic and mutagenic DNA double-strand breaks (2, 3). Consequently, cellular processes regulated in an ATM-dependent manner include chromatin decondensation, apoptosis, senescence, cell cycle, redox balance, metabolism, and splicing (4).

A-T has a wide range of clinical manifestations, however, one of the most devastating clinical signs of the classical form of A-T is neurodegeneration. The disease manifests as motor dysfunction in young children and is predominantly characterised by the progressive loss of Purkinje and granule neurons in the cerebellum (5). Cerebellar degeneration is thought to be linked with defects in the neuronal DNA damage response, metabolic abnormalities, and epigenetic silencing of diverse neuronal genes (4, 5). However, the molecular mechanisms that underpin cerebellar degeneration in A-T are poorly understood.

Microglia are resident immune cells of the central nervous system (CNS). Microglia play instrumental roles in the development and maintenance of the CNS, including regulation of neurogenesis, synaptic maintenance and plasticity, trophic support of other cell types, and clearance of apoptotic and dead cells. However, persistent microglial activation and consequent neuroinflammation are implicated in the pathology of diverse neurodegenerative disorders (6). For example, somatic mutations in the BRAF oncogene in erythro-myeloid precursors of microglia result in progressive neurological impairment with multiple features of cerebellar ataxia (7). In addition, microglial priming (an enhanced response to secondary stimuli) and activation, which are both linked to neuroinflammation, are seen in mouse models of *Ercc1^Δ/-^-linked* nucleotide excision repair and frataxin deficiencies (8, 9).

In A-T defective mice and rats morphological changes associated with microglial activation have been observed (10–12). Pharmacological inhibition of Atm in cultured *ex vivo* murine microglia led to neuronal cell death, which was mediated by the secretion of the neurotoxic cytokine IL-1β (12). It was suggested that microglia mount a neurotoxic innate immune response to cytosolic DNA (10–12). In addition, links between ATM deficiency and activation of the innate immune signalling have been demonstrated in other cell types (13–15). Rodent models of ATM deficiency, however, do not fully recapitulate the human phenotype, possibly due to a higher sensitivity of the human CNS to oxidative stress and DNA damage (11, 16). Moreover, microglia from rodents and humans show fundamental differences in their proliferation, response to extracellular signalling molecules and secretion, and exhibit highly divergent gene expression signatures (17–19).

In the present work, we utilised human cell models to investigate the underlying basis for the role of microglia in neurodegeneration in A-T. We find that loss of ATM, or its activity, promotes sustained microglial activation linked with increased expression of pro-inflammatory cytokines and phagocytic clearance. Microglial activation was shown to be mediated by the non-canonical RELB/p52 nuclear factor NF-κB transcriptional pathway. The RELB/p52 pathway is activated in response to persistent DNA damage associated with A-T and is driven by the NF-κB-inducing NIK kinase. Chronic activation of ATM-deficient microglia results in excessive phagocytosis of neurites, potentially contributing to neurodegeneration. These data provide mechanistic insights into microglial dysfunction in human A-T.

## RESULTS

### Loss of ATM function results in microglial activation

The effects of ATM deficiency on microglial function were studied in human HMC3 and C20 microglial cell lines (20–22). To provide an *ATM*-deficient model system, that is representative of A-T (23), an *ATM* knockout (KO) human microglial HMC3 cell line was generated using CRISPR/Cas9 (Supplementary Figure S1A and S1B). The loss of ATM function was verified by treating *ATM* KO microglia with camptothecin, a topoisomerase I inhibitor that induces ATM activation (24). As expected, *ATM* KO microglia displayed: i) no visible autophosphorylation of ATM at S1981 and grossly attenuated phosphorylation of ATM’s downstream target, CHK2, in response to camptothecin (Figure 1A) (25), and ii) reduced cell proliferation compared to wild-type (WT) cells (Supplementary Figure S1C). To determine whether the *ATM* KO microglia were activated, we first measured expression levels of two markers of microglial activation, CD40 and CD68 (26, 27). Cell surface levels of CD40 were increased in *ATM* KO microglia to a similar extent as in WT cells stimulated with TNFα (Figure 1B). Additionally, the levels of lysosomal CD68 were higher in *ATM* KO microglia as compared to WT (Supplementary Figure S1D). To further investigate the activation status of *ATM* KO microglia, we measured mRNA expression of the pro-inflammatory cytokines *IL6*, *IL8*, *IL1B* and *TNFA*. All were up-regulated by 3-12-fold in *ATM* KO *vs* WT cells, whereas the mRNA levels of anti-inflammatory *IL4* were unaffected by ATM status (Figure 1C). The microglial activation phenotypes were rescued upon re-expression of ATM from a doxycycline-inducible piggyBac-transgene (28), showing that the effects were specific for ATM loss (Figure 1A-C; Supplementary Figure S1E).

**Figure 1:**
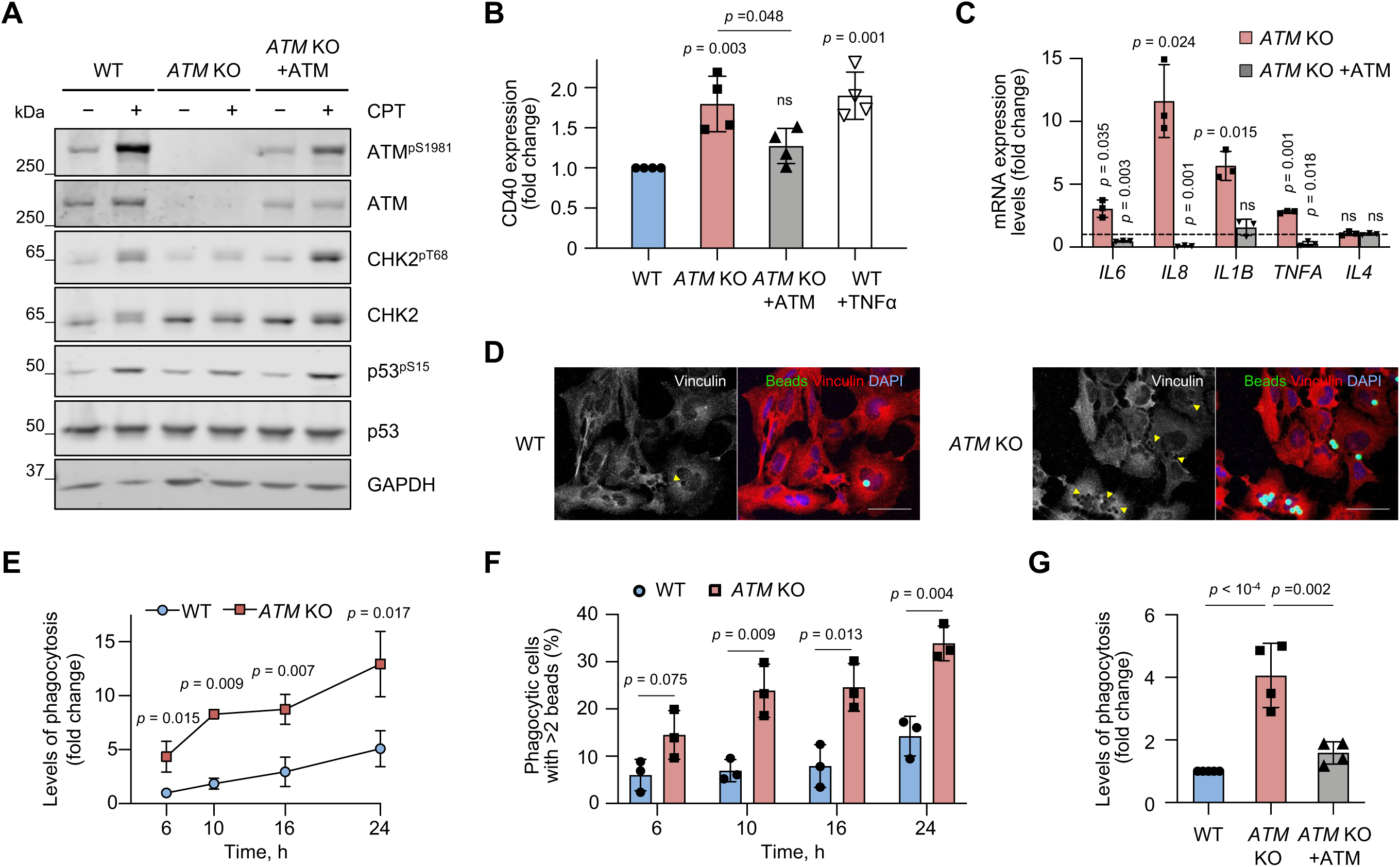
ATM-knockout (KO) microglia are activated, resulting in increased expression of pro-inflammatory cytokines and enhanced phagocytic clearance. **A** Immunoblot analysis of WT, *ATM* KO and *ATM* KO cells in which ATM was re-expressed from a piggyBac transgene. To test for ATM activity, cells were treated with 1 μM camptothecin (CPT) for 1 h. Loading control: GAPDH. **B** Protein expression levels of the microglial activation marker, CD40, measured by flow cytometry in cells as in (**A**). Expression is relative to WT HMC3. Positive control: 50 µg/mL TNFα for 4 h, cells collected for analysis at 20 h post-treatment. Mean ± S.D. shown (n = 4). **C** Quantitative RT-qPCR analysis of the indicated cytokines in cells as in (**A**). Expression is relative to WT HMC3 (dashed line). Reference genes: *RSP13* and *IPO8*. Mean ± S.D. shown (n = 3). One-sample t-test used. **D** Representative immunofluorescence images of phagocytic uptake of 5 μm beads in WT and *ATM* KO HMC3 (6 h assay). Confocal Z-stack compression images shown. Arrowheads indicate disruption in vinculin structure associated with bead engulfment. Scale bar: 50 µm. Bead substrates (green), vinculin (red), DNA (blue). **E** Kinetics of phagocytosis (5 μm beads) in WT and *ATM* KO cells at the indicated time points. Phagocytosis is relative to WT HMC3 at 6 h with 3.7 ± 1.3% of phagocytic cells. Mean ± S.D. shown (n = 3). **F** Kinetics of the changes in percentage of highly phagocytic cells (> 2 beads per cell) as in (**E**). To determine the number of beads per cell, Mean Fluorescence Intensity (MFI) of the population of phagocytic cells was divided by MFI of a single 5 µm fluorescent bead. Mean ± S.D. shown (n = 3). Unpaired two-tailed t-test used. **G** Phagocytosis levels (6 h assay) of cells as in (**A**). Phagocytosis is relative to WT HMC3, in which 5.8 ± 3.2% of cells are phagocytic. Mean ± S.D. shown (n = 4). **B, E, G** One-way ANOVA with Tukey’s multiple comparison’s test used.

One indicator of microglial activity is their ability to engulf cell debris, apoptotic and stressed cells, which expose the “eat-me” signal, phosphatidylserine, on their surface (6). Phagocytic properties of WT and *ATM* KO microglia were investigated using 5 µm carboxylated latex beads, which mimic the size of neuronal soma with externalised phosphatidylserine, as substrates (Supplementary Figure S2A). Both the percentage of phagocytic cells, further referred to as levels of phagocytosis, and phagocytic activity, which reflects the relative fluorescence intensity of engulfed beads per cell, were measured using flow cytometry (Supplementary Figure S2B). We observed an increase in: i) the percentage of phagocytic *ATM* KO cells compared to WT, over a 24 h time period, using immunofluorescence and flow cytometry (Figure 1D and 1E), and ii) the fraction of highly phagocytic *ATM* KO cells, which engulfed more than 2 beads per cell, at different time points compared with WT (Figure 1F). The kinetics of phagocytic changes was linear in both cell lines over a 24 h period. However, to exclude any early apoptotic changes due to uptake of non-digestible substrates, the assays were carried out for 6 h. Also, given that the changes in phagocytic activities were modest, possibly due to the relatively large size of the substrates, levels of phagocytosis were used as the primary readout in subsequent assays. Upon re-expression of ATM, enhanced phagocytic properties of *ATM* KO cells were rescued (Figure 1G; Supplementary Figure S2C). Similar to *ATM* KO microglia, increased levels of phagocytosis were observed in HMC3 and C20 human microglia, in which ATM was knocked down using siRNA (Supplementary Figure S2D-G). These data indicate that ATM-deficient microglia are more efficient phagocytes than their WT counterparts, and this effect is specific to ATM loss and cell line independent.

To investigate whether enhanced phagocytic properties of ATM-deficient microglia are mediated via the loss of ATM protein or its kinase activity, WT cells were treated with a reversible ATM inhibitor, AZD1390 (29), for up to 9 days (Supplementary Figure S3A and S3B). To rescue the effects of ATM inhibition, AZD1390 was washed out and the cells were allowed to recover for 2 days (Supplementary Figures S3A and S3B, Release). The levels of ATM autophosphorylation and CHK2 phosphorylation following treatment with camptothecin indicated that kinase inhibition and inhibitor wash-out were successful (Supplementary Figure S3A). We then determined phagocytosis levels of AZD1390-treated WT cells and respective DMSO-treated controls. Although no significant changes in phagocytosis levels were observed at day 1 of inhibitor treatment, we found a 2-2.5-fold increase at days 4-6 in the ATM-inhibited cells compared to the control (Supplementary Figure S3B). It is likely that the long-term effects of ATM loss/inhibition, such as the accumulation of oxidative stress and DNA damage, drive the changes in phagocytic properties. Importantly, inhibitor removal resulted in a reduction in phagocytosis in ATM inhibitor-treated cells back to the levels of DMSO-treated microglia (Supplementary Figure S3B, Release).

Together, these data indicate that loss of ATM results in persistent microglial activation, which is characterised by increased expression of pro-inflammatory cytokines and enhanced ability to engulf synthetic substrates. These phenotypes are specific to ATM and driven by long-term changes associated with loss of ATM and/or its kinase activity.

### The non-canonical RELB/p52 NF-κB pathway is activated in ATM-deficient microglia

The NF-κB proteins are central mediators of inflammation in response to tissue damage and infection (30). If mis-regulated, normally protective NF-κB-mediated pro-inflammatory responses can amplify acute or chronic tissue damage, thus driving autoinflammatory and autoimmune disease (31). NF-κB family members include RELA (p65), RELB, c-REL, p50 and p52. The canonical NF-κB pathway provides a rapid and transient response to cytokines, mitogens, and growth factors. These stimuli activate the IκB kinase (IKK) complex, which phosphorylates IκBα, inducing its proteasomal degradation. Loss of inhibitory binding of IκBα to NF-κB proteins results in nuclear translocation of the canonical NF-κB dimers, the most abundant of which are p65-p50 and c-REL-p50. The precursor protein p105 also acts in the NF-κB-inhibitory manner and becomes active upon proteolytic processing into p50 upon stimulation (31).

Microglial activation and neuroinflammation are linked to the activities of the p65-p50 heterodimer (32). In addition, p65 is localised to the nucleus in rodent models of A-T (11, 12). We therefore investigated whether the canonical NF-κB pathway is activated in ATM-deficient human microglia. We first tested whether canonical NF-κB signalling can be induced in WT microglia upon treatment with the tumour necrosis factor TNFα, a well-established activator of the pathway (31). We observed TNFα-dependent: i) phosphorylation of p65 at S536 indicative of its activation (33) and relocalisation of p65 in the nucleus (Figure 2A and 2B; Supplementary Figure S4A and S4B), and ii) processing of p105 to p50 and nuclear translocation of p50 in WT cells (Figure 2B). These data indicate that the canonical NF-κB pathway is inducible in WT microglia. In contrast, while we observed a modest increase in: i) basal p65 phosphorylation at S536 and ii) processing of p105 into p50 in *ATM* KO cells as compared to WT, we failed to detect relocalisation of p65 and p50 into the nucleus (Figure 2A and 2B; Supplementary Figure S4A and S4B). A similar lack of nuclear translocation of p65 was seen in *ATM* KO C20 microglia, confirming that the effect was not cell-line specific (Supplementary Figure S4C-E). To investigate the reasons of this effect, protein levels of the NF-κB-inhibitory protein IκBα were determined. We found that IκBα levels were only moderately reduced in ATM-deficient microglia as compared to WT cells (Figure 2A), and propose that the remaining IκBα is sufficient to retain the p65-containing complexes in the cytoplasm.

**Figure 2:**
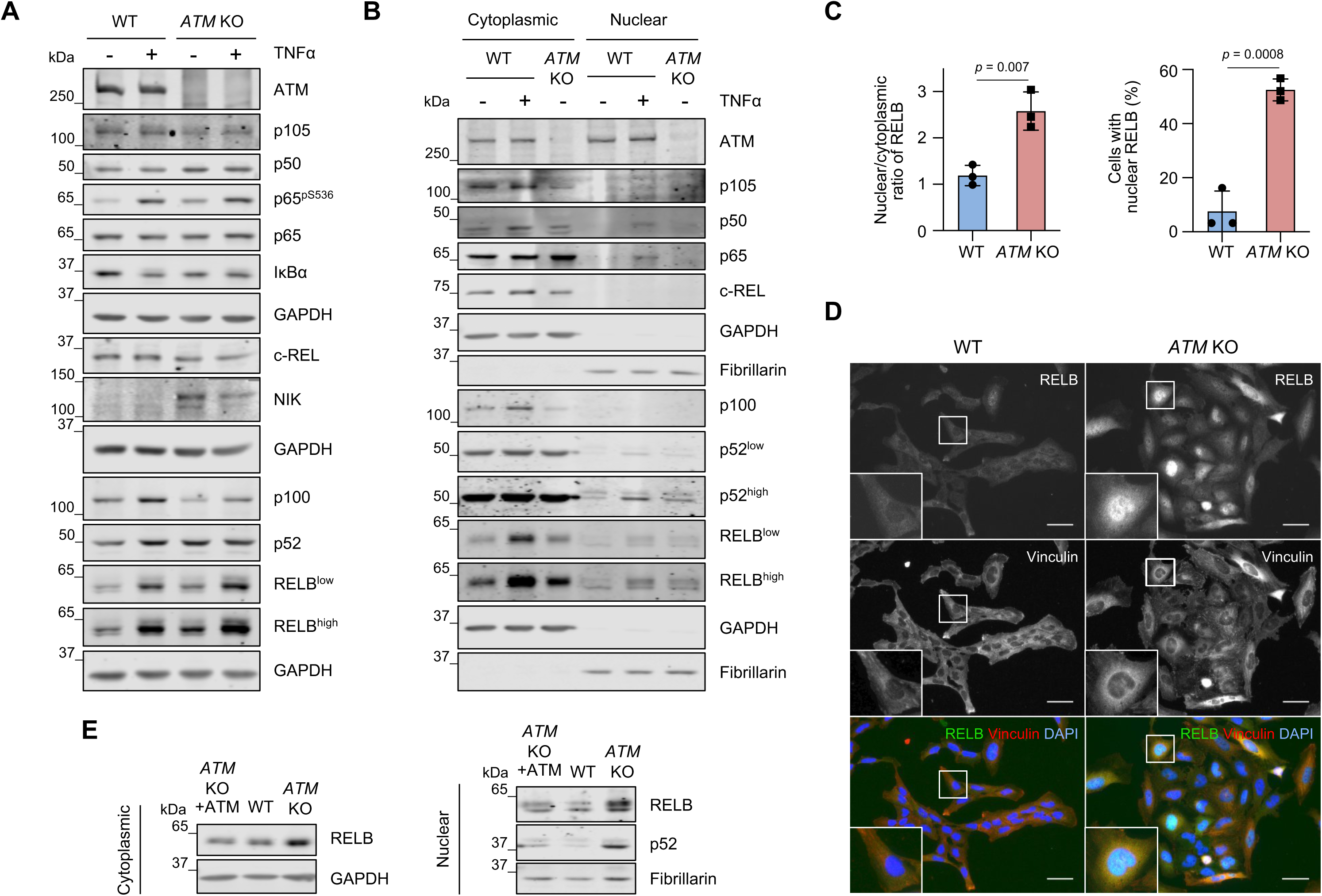
The RELB/p52 non-canonical NF-κB pathway is activated in ATM-deficient microglia. **A** Immunoblot analysis of NF-κB protein levels in WT and *ATM* KO HMC3 microglia using the indicated antibodies. Positive control: 50 µg/mL TNF-α for 6 h. Loading control: GAPDH. **B** Immunoblot analysis of nuclear relocalisation of NF-κB family members in WT and *ATM* KO HMC3 as in (**A**). Loading controls: GAPDH (cytoplasmic), fibrillarin (nuclear). **C** Quantification of RELB localisation presented as nuclear/cytoplasmic ratio (relative to WT; left) and percentage of cells with nuclear RELB (right) in WT and *ATM* KO HMC3 cells. Mean ± S.D. shown (n = 3). Unpaired t-test used. **D** Representative images of (**C**). Images in a single Z-plane shown. Scale bar: 50 µm. RELB (green), vinculin (red), DNA (blue). **E** Immunoblot analysis of RELB levels and localisation in cytoplasmic and nuclear extracts of WT, *ATM* KO and *ATM* KO HMC3, in which ATM was re-expressed as in Figure 1A. Loading controls: GAPDH (cytoplasmic), fibrillarin (nuclear).

Additionally, as c-REL-containing heterodimers play a role in the canonical NF-κB response, the cellular localisation of c-REL was determined in WT and *ATM* KO HMC3 microglia (31). Although we failed to observe nuclear translocation of c-REL following TNFα treatment (Figure 2A and 2B), HMC3 cells were previously shown to have proficient c-REL signalling (34). Similarly to p65, c-REL was retained in the cytoplasm, confirming that the c-REL-dependent NF-κB pathway is not activated in ATM-deficient microglia (Figure 2B). Together, these data indicate that proteasomal degradation of IκBα and the nuclear translocation of p65, p50 and c-REL serve as limiting factors to canonical NF-κB activation in the absence of ATM.

Non-canonical NF-κB signalling is activated downstream of cell surface receptors of the tumour necrosis factor receptor (TNFR) superfamily, such as BAFFR, CD40, LTβR, and RANK. Pathway activation is dependent on stabilisation of the NIK (NF-κB-inducing) kinase, which phosphorylates IKKα, promoting IKKα-mediated phosphorylation of p100 and its cleavage-dependent processing to p52. RELB-p52 is the major non-canonical heterodimer that drives transcriptional programmes (35).

To investigate whether the non-canonical NF-κB pathway is activated in ATM-deficient microglia, we first determined the levels of NIK and the efficiency of p100 processing by immunoblotting. NIK protein levels were increased in the *ATM* KO cells, compared to WT, indicating its enhanced stability (Figure 2A). In addition, a reduction in basal levels of p100 and a concurrent increase in p52 levels were observed in *ATM* KO microglia compared to the WT control, indicating enhanced p100 processing (Figure 2A and 2B). We also determined the basal levels of RELB and its cellular localisation in WT and *ATM* KO HMC3 cells (Figure 2A-E; Supplementary Figure S5A). The specificity of the antibody against RELB, which also detects various post-translationally modified forms of the protein (33), was confirmed using siRNA-mediated knockdown (Supplementary Figure S5A). In agreement with the *de novo* synthesis of RELB prior to nuclear translocation (35), basal RELB levels were increased in the absence of ATM compared to the WT control (Figure 2A and 2B; Supplementary Figure S5A). Additionally, RELB was detected in the nucleus of *ATM* KO cells whereas almost no nuclear RELB was found in the WT control (Figure 2B; Supplementary Figure S5A). These findings were independently confirmed using immunofluorescence. An increase in the nuclear to cytoplasmic (N/C) ratio and in the percentage of cells with nuclear RELB was observed in *ATM* KO cells *vs* the WT control (Figure 2C and 2D). Noteworthy, the increased levels and nuclear translocation of RELB and p52 observed in ATM *KO* cells were rescued upon re-expression of ATM (Figure 2E). Finally, a similar basal increase in nuclear RELB was seen in *ATM* KO C20 microglia (Supplementary Figures S5B and S5C).

To confirm and extend these results, RELB levels and localisation were determined in WT cells treated with the ATM kinase inhibitor, AZD1390, for up to 9 days (Supplementary Figure S3C). A gradual increase in RELB expression and nuclear translocation was observed at days 3-6 of treatment, reaching a plateau at days 6-9. No changes were seen in the DMSO-treated control (Supplementary Figure S5D). Together, these results unequivocally show that loss of ATM function results in specific activation of the RELB/p52 non-canonical NF-κB pathway in human microglia.

### The non-canonical NF-κB pathway controls microglial activation in the absence of ATM

Having observed the stabilisation of NIK kinase and nuclear translocation of RELB, but not p65 or C-REL, in the absence of ATM, we aimed to determine whether RELB-containing complexes might be responsible for microglial activation. Therefore, the phagocytic properties and expression levels of several pro- and anti-inflammatory cytokines were investigated in WT and *ATM* KO microglia following siRNA-mediated knockdown of RELB. In addition, siRNA that targets all five NF-κB subunits (NF-κB^pan^) was used to probe for synergistic effects of the canonical and non-canonical pathways (Figure 3A-C). The knockdown of RELB significantly reduced phagocytosis levels in *ATM* KO cells to levels typical of WT cells. No effect of RELB loss on phagocytic properties of WT cells was observed, indicating that the phagocytic properties of ATM-deficient microglia are modulated via RELB (Figure 3B).

**Figure 3:**
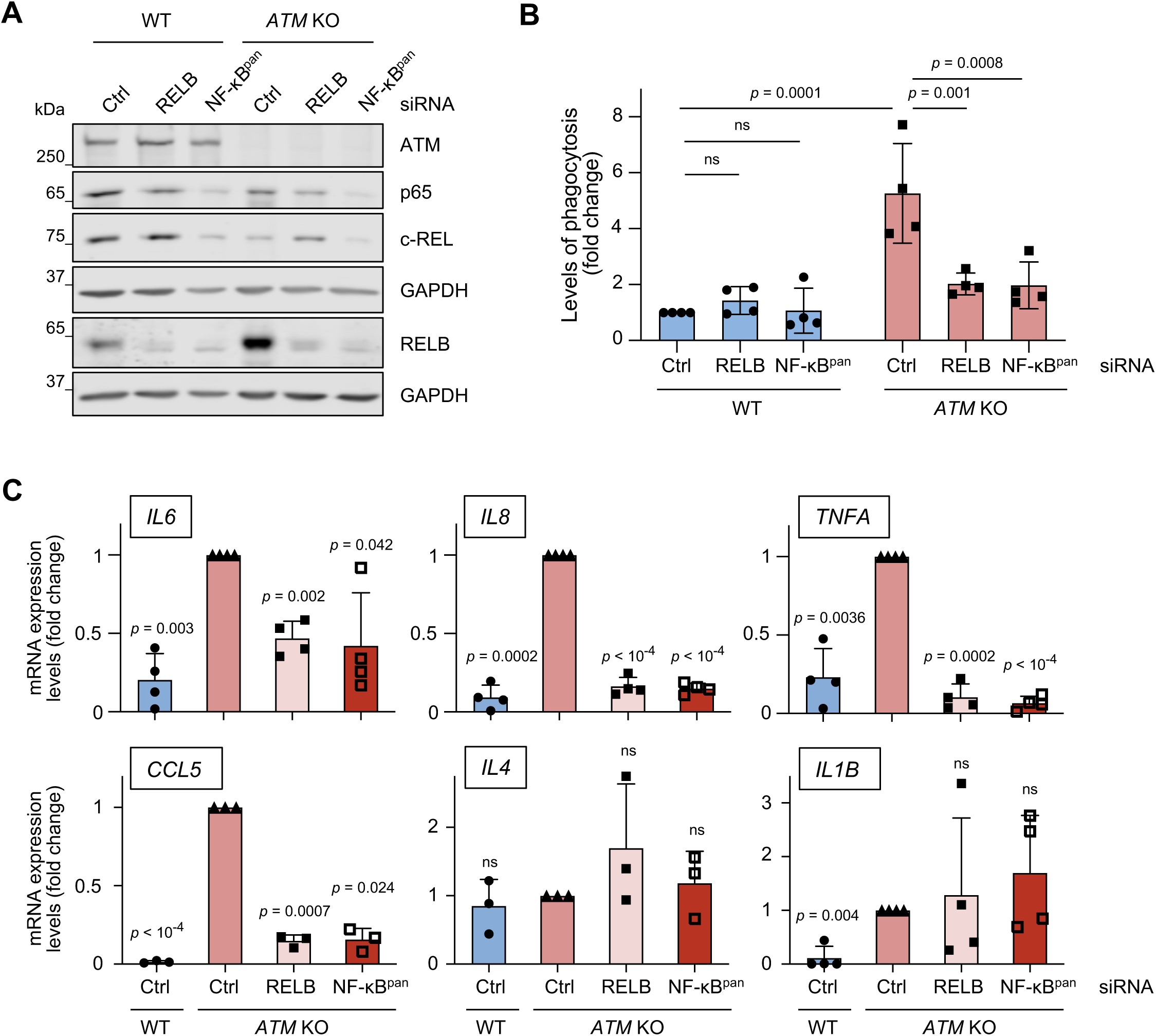
Microglial activation in the absence of ATM is mediated via RELB. **A** Immunoblot analysis of siRNA-mediated silencing of RELB or all NF-κB members (NF-κB^pan^) in WT and *ATM* KO HMC3 cells. Loading control: GAPDH. **B** Phagocytosis levels (5 μm beads) of WT and *ATM* KO HMC3 cells treated with control (Ctrl), RELB or NF-κB^pan^ siRNA. Phagocytosis is relative to WT HMC3 treated with Ctrl siRNA. Mean ± S.D. shown (n = 4). Two-way ANOVA with Tukey’s multiple comparison’s test used. **C** Quantitative RT-qPCR analysis of the indicated cytokines as in (**B**). Expression is relative to *ATM* KO HMC3 treated with control siRNA (Ctrl). Reference gene: *RSP13*. Mean ± S.D. shown (*IL6, IL8, TNFA, IL1B* n = 4, *IL4, CCL5* n = 3). One sample t-test used.

Next, we studied the effect of RELB depletion on cytokine expression. As expected, siRNA transfection moderately enhanced the fold change difference in mRNA levels of pro-inflammatory cytokines *IL6*, *IL8*, *TNFA, CCL5* and *IL1B* but not anti-inflammatory *IL4* in *ATM* KO *vs* WT control, as compared to that seen in unchallenged cells (Figure 3C, compare Ctrl siRNA-treated *ATM* KO *vs* WT cells with the results in Figure 1C). Importantly, upon RELB knockdown, the high expression levels of NF-κB-dependent cytokines *IL6*, *IL8* and *TNFA* in Ctrl-siRNA treated *ATM* KO microglia was reduced to similar levels as observed in Ctrl-siRNA treated WT cells (Figure 3C). Expression of the chemokine *CCL5* was reduced in the absence of ATM and RELB but to a lesser extent, in agreement with it being cooperatively regulated by NF-κB and interferon-regulatory factors IRF1, IRF3 and IRF7 (36). In contrast, mRNA expression levels of *IL1B* that acts upstream of NF-κB (37), and anti-inflammatory *IL-4* (38), remained unchanged in *ATM* KO cells treated with RELB siRNA (Figure 3C). One important observation is that both phagocytic properties and expression of pro-inflammatory cytokines were reduced at similar levels following treatment of ATM-deficient microglia with siRNA against RELB and all NF-κB subunits, confirming the dependency of these phenotypes on RELB-containing complexes (Figure 3B and 3C).

These data demonstrate that the RELB/p52 non-canonical NF-κB pathway promotes microglial activation in the form of enhanced phagocytic clearance and expression of pro-inflammatory cytokines in the absence of ATM. We therefore further investigated the mechanisms of RELB/p52 activation in ATM-deficient microglia.

### RELB/p52-dependent activation of ATM-deficient microglia is mediated by NIK kinase

To determine whether enhanced RELB/p52 signalling in the absence of ATM is dependent on NIK kinase, we used siRNA to knock down NIK and investigated the microglial activation phenotypes (Figure 4A-D). As expected from the dependence of processing of p100 to p52 on the NIK-IKKα axis (35), NIK downregulation resulted in the modest accumulation of p100 and consequent reduction in p52 protein levels in both WT and *ATM* KO cells (Figure 4A and 4B). Importantly, NIK loss partially rescued: i) the nuclear translocation of RELB (Figure 4A), ii) the expression of NF-κB-dependent pro-inflammatory cytokines *IL8, TNFA* and *CCL5*, whereas there was no effect on the levels of anti-inflammatory *IL4* (Figure 4C), and iii) the levels of phagocytosis (Figure 4D) in *ATM KO*, and not WT microglia. These data show that RELB/p52 non-canonical NF-κB signalling and microglial activation in the absence of ATM are dependent on NIK kinase and are likely to be regulated, at least in part, via extracellular signalling cues.

**Figure 4:**
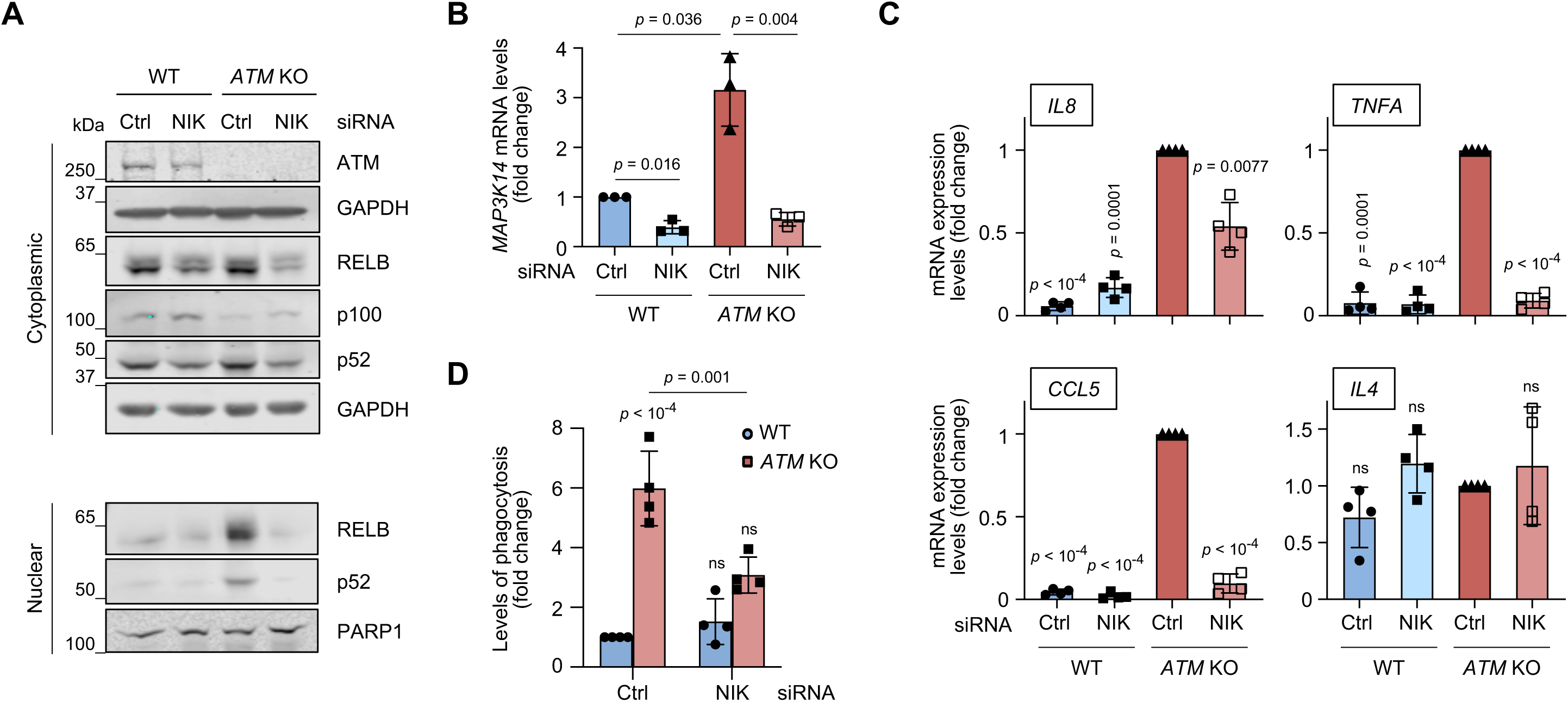
Microglial activation and non-canonical NF-κB signalling in ATM-deficient cells are mediated by NIK kinase. **A** Immunoblot analysis of the non-canonical NF-κB pathway proteins in cytoplasmic and nuclear extracts of NIK-deficient WT and *ATM* KO HMC3 microglia. Loading controls: GAPDH (cytoplasmic), PARP1 (nuclear). **B** Quantitative RT-qPCR analysis of NIK knockdown efficiency in WT and *ATM* KO HMC3 microglia. Expression is relative to Ctrl siRNA-treated WT HMC3. Reference gene: *RSP13*. Mean ± S.D. shown (n = 3). Two-way ANOVA with Tukey’s multiple comparison’s test was used for 2^-ΔCt^ values. **C** Quantitative RT-qPCR analysis of the indicated cytokines as in (**A, B**). Expression is relative to Ctrl siRNA-treated *ATM* KO HMC3. Reference gene: *RSP13*. Mean ± S.D. shown (n = 4). One sample t-test used. **D** Phagocytosis levels (5 μm beads) of cells treated as in (**A, B**). Phagocytosis is relative to Ctrl siRNA-treated WT HMC3, in which 3.5 ± 1.7% of cells are phagocytic. Mean ± S.D. shown (n = 4). Two-way ANOVA with Tukey’s multiple comparison’s test used.

### Persistent DNA damage promotes RELB/p52 NF-κB signalling and microglial activation

The continuous occurrence of DNA damage requires proficient DNA damage signalling and repair, including ATM-dependent responses, to maintain genome stability. A-T cells are therefore characterised by enhanced oxidative stress and delayed repair of a subset of specialised DNA lesions (2,3,39). Importantly, DNA damage is thought to activate the non-canonical NF-κB pathway, although the mechanism of this activation is unclear (30). We therefore set out to investigate whether DNA damage might drive RELB/p52-dependent activation of microglia.

To determine whether *ATM* KO microglia possess increased levels of spontaneous DNA damage, we analysed the cells for several DNA damage markers. Protein poly[ADP-ribosyl]ation (PAR) and phosphorylation of histone variant H2AX at S139 (γH2AX), prototypical signalling events that occur during the DNA damage response (4, 40), were increased in ATM-deficient cells (Figure 5A). Additionally, the basal levels of DNA damage were directly measured using alkaline single-cell gel electrophoresis, which detects alkali labile sites, DNA single- and double-strand breaks (41). Basal levels of DNA damage were elevated in *ATM* KO microglia compared to the WT control (Figure 5B). We also analysed the levels of intracellular reactive oxygen species (ROS), which serve as one of the sources of DNA damage in ATM-deficient cells. Increased intracellular ROS were detected in *ATM* KO microglia, in comparison with WT (Figure 5C).

**Figure 5:**
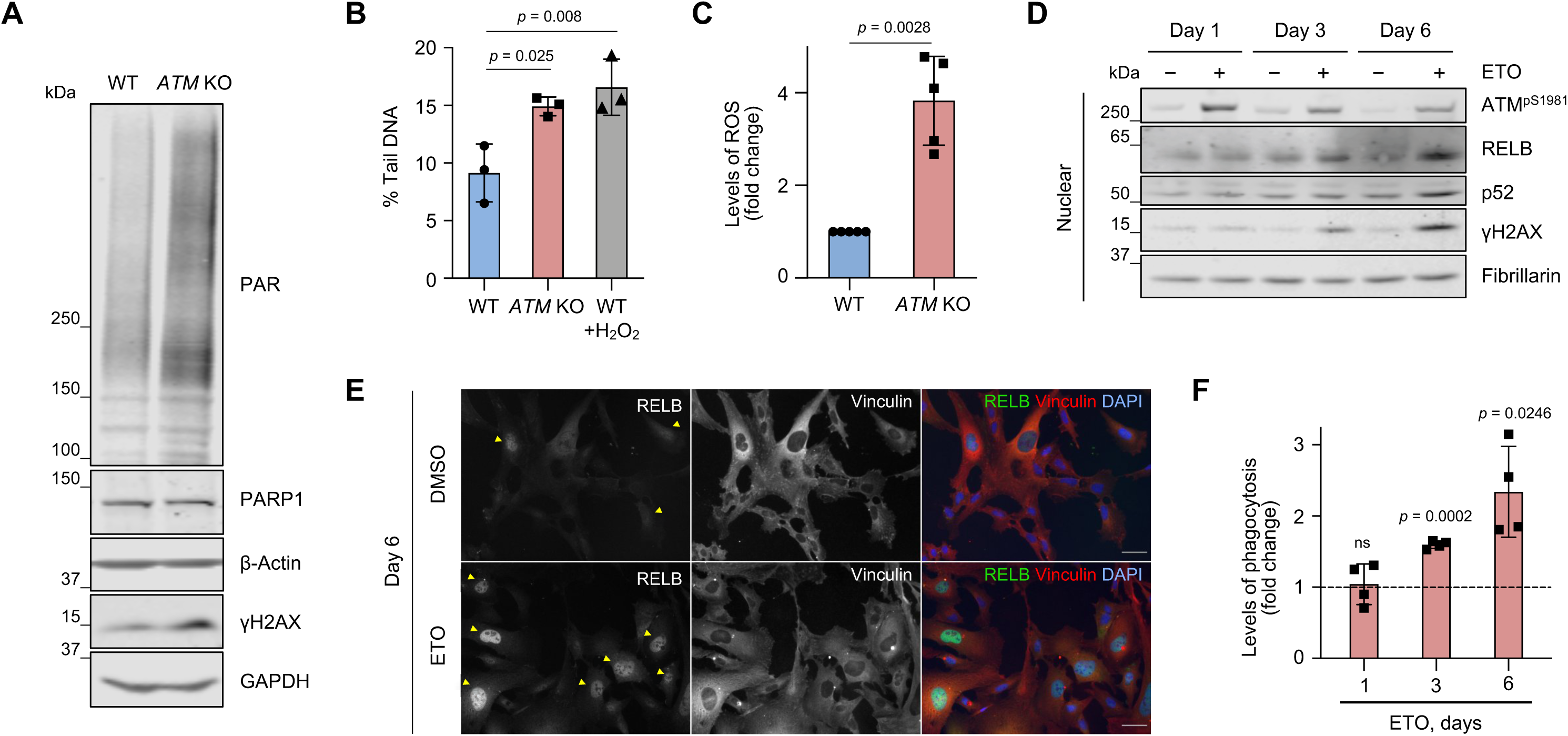
Persistent DNA damage promotes microglial activation and non-canonical NF-κB signalling. **A** Immunoblot analysis of DNA damage response markers in WT and *ATM* KO HMC3 cells using the indicated antibodies. Loading controls: β-actin, GAPDH. **B** Levels of endogenous DNA damage in WT and *ATM* KO HMC3 detected by alkaline Comet assays and presented as % tail DNA. Positive control: 2 µM H_2_O_2_ for 5 min. Mean ± S.D. shown (technical duplicates per experiment, n = 3). One-way ANOVA with Dunnett’s multiple comparison’s test used. **C** Levels of intracellular reactive oxygen species (ROS) in WT and *ATM* KO HMC3 measured using DCF-DA probe and flow cytometry. ROS levels are relative to WT HMC3. Mean ± S.D. shown (n = 5). One sample t-test used. **D** Immunoblot analysis of DNA damage response markers and RELB/p52 levels in nuclear fractions of WT HMC3 microglia treated with 0.5 μM etoposide (ETO) for up to 6 days. Control: DMSO. Loading control: fibrillarin. **E** Representative images of cells treated as in (**D**). Images in a single Z-plane shown. Arrowheads indicate cells with nuclear RELB. Scale bar: 50 µm. RELB (green), vinculin (red), DNA (blue). **F** Phagocytosis levels (5 μm beads) of WT microglia treated as in (**D**). Phagocytosis is relative to DMSO control at respective time points (dashed line). Mean ± S.D. shown (n = 3). One sample t-test used.

To induce persistent low-level DNA damage, as seen in ATM-deficient microglia, WT HMC3 cells were treated with a low dose of etoposide for 6 days, a topoisomerase II inhibitor that induces DNA double-strand breaks (42) (Figure 5D-F; Supplementary Figure S6A-C). The presence of DNA damage was confirmed by the increased levels of γH2AX and autophosphorylation of ATM at S1981 (Figure 5D; Supplementary Figure S6A). The concentration of etoposide was sufficiently low to ensure that no visible apoptosis and/or necrosis was observed following 6 days of continuous treatment, as adjudged by the lack of cleaved caspase-3 positive staining or condensed and fragmented nuclei (Supplementary Figure S6A). Importantly, the treatment resulted in a time-dependent increase in: i) the levels of nuclear RELB and p52 in the absence of p65 translocation in the nucleus (Figure 5D and 5E; Supplementary Figure S6B and S6C) and ii) the levels of phagocytosis as compared to respective DMSO controls (Figure 5F). The kinetics of DNA damage-induced changes in nuclear translocation of RELB and phagocytosis was similar to that observed in response to long-term treatment with AZD1390 (Supplementary Figure S3B and S5D). These results indicate that persistent low-level DNA damage associated with ATM loss underpins microglial activation mediated by the non-canonical RELB/p52 pathway.

### ATM-deficient microglia excessively engulf neurites

In addition to secretion of pro-inflammatory cytokines and oxidative species, chronically activated microglia can contribute to neurodegenerative pathologies by: i) defective phagocytosis of dead and dying cells that leads to uncontrolled release of neurotoxic compounds and ii) excessive phagocytosis of neuronal synapses, axons and dendrites (43–46). To investigate the effects of microglial ATM deficiency on neuronal engulfment, we employed post-mitotic dopaminergic neurons obtained by differentiation of human mesencephalic LUHMES progenitors (47) (Supplementary Figure S7A and S7B). To model ATM’s loss-of-function, post-mitotic neurons were maintained in the presence of AZD1390 during differentiation days 2-5. After AZD1390 was washed out, ATM activity remained inhibited for at least 48 h, during which time the ATM-deficient neurons were used in assays (Supplementary Figure S7C). The 3-day inhibition of ATM did not induce measurable neuronal apoptosis in post-mitotic LUHMES (Supplementary Figure S7D). In addition, we observed no damaging effect of factors secreted by WT or *ATM* KO microglia on ATM-deficient neurons (Supplementary Figure S7E and S7F).

For the phagocytosis assays, microglia and neurons were labelled with the fluorescent live dyes carboxyfluorescein succinimidyl ester (CFSE) and CellTrace Violet (CTV), respectively, and cultured together for 24 h (Figure 6A). The presence of CTV-positive neuronal material within CFSE-positive microglia (double positive cells) was assessed by flow cytometry. ATM *KO* microglia consistently engulfed (50 ± 13)% more ATM-deficient neuronal material than WT microglia (Figure 6B, left). Unexpectedly, a similar increase in the engulfment of healthy WT neurons by *ATM* KO microglia, as compared with WT, was observed (Figure 6B, right). These results indicate that microglial phagocytosis in the absence of ATM is regulated via cell-intrinsic mechanisms.

**Figure 6:**
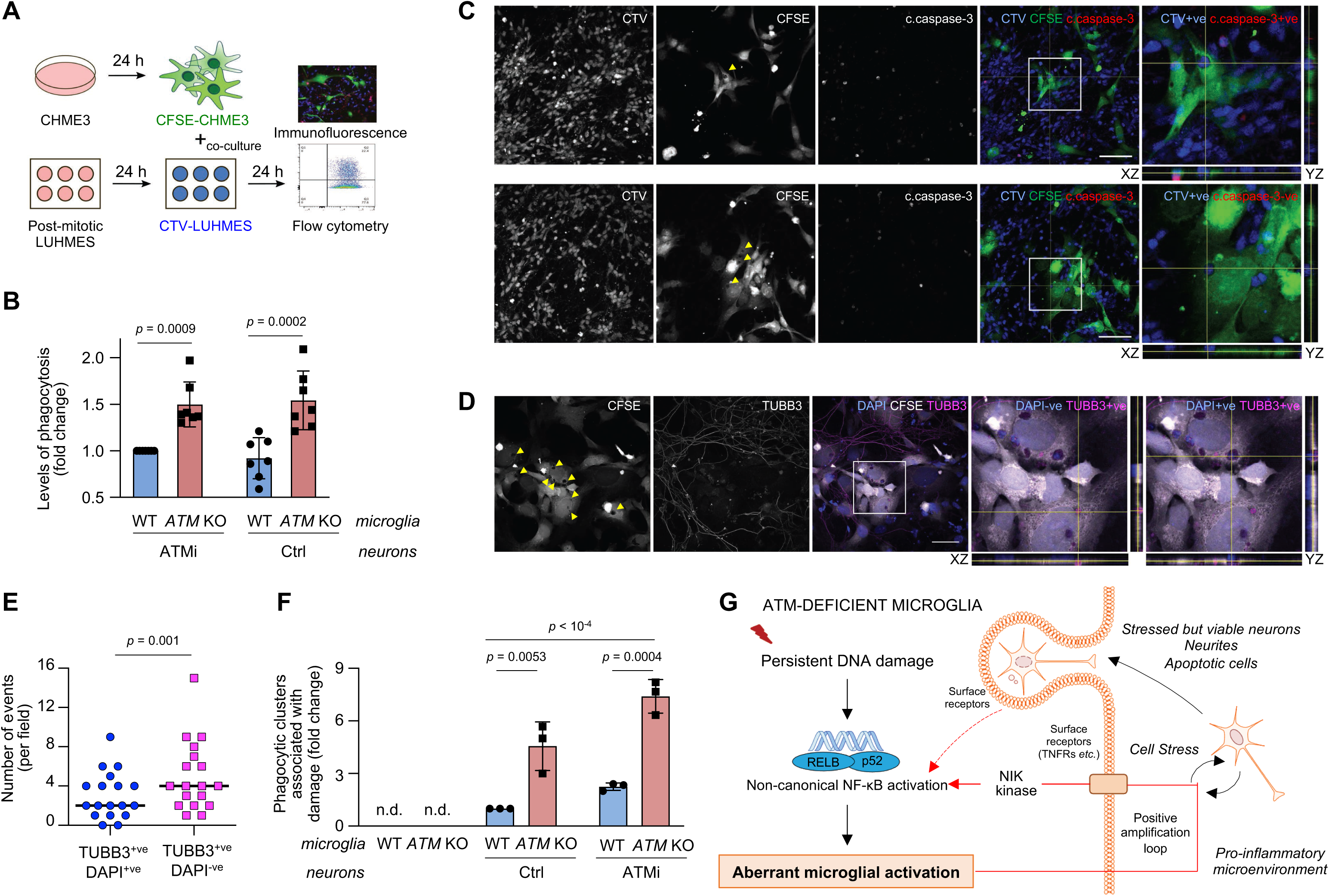
Neurotoxicity of ATM-deficient microglia is linked with aberrant phagocytosis of healthy neuronal soma and neurites. **A** Schematic of neuronal-microglial co-culture assays. Phagocytic uptake of LUHMES neurons by HMC3 microglia was analysed by flow cytometry and immunofluorescence. **B** Phagocytosis levels (CTV-stained post-mitotic LUHMES) of WT and *ATM* KO CFSE-labelled HMC3 cells using post-mitotic LUHMES neurons. LUHMES cells were pre-treated with AZD1390 for 3 days and released from inhibition prior to setting up the co-cultures (ATMi) or left untreated (Ctrl). Phagocytosis is relative to WT HMC3/ATMi LUHMES. Mean ± S.D. shown (n = 4). Two-way ANOVA with Sidak’s multiple comparisons used. **C, D** Representative immunofluorescence images of phagocytosis events by *ATM* KO microglia as in (**B**). Confocal Z-stack compression images and relevant XY and YZ projections shown. Scale bar: 50 μm. **C** Arrows indicate microglial phagolysosomes containing engulfed neuronal bodies (CTV+ve), which are positive (top) or negative for cleaved caspase-3 (bottom). CFSE-labelled *ATM* KO HMC3 microglia (CFSE; green), cleaved caspase-3 (c.caspase-3; red), CTV-labelled LUHMES neurons (CTV; blue). **D** Arrows indicate microglial phagolysosomes containing neuronal soma (TUBB3+ve, DAPI+ve) and neurites (TUBB3+ve, DAPI-ve). CFSE-labelled *ATM* KO HMC3 microglia (CFSE; green), β3-tubulin (TUBB3; red), DAPI (blue). **E** Quantification of phagocytic events as in (**D**). Distribution of TUBB3+ve DAPI+ve *vs* TUBB3+ve DAPI-ve events per field of view shown (n = 20 fields of view from two independent biological experiments). Wilcoxon matched-pairs signed rank test used. **F** Quantification of phagocytic microglial clusters associated with damage to the neuronal network as in (**B**). Data are relative WT microglia co-cultured with untreated LUHMES post-mitotic neurons (Ctrl). Mean ± S.D. shown (n = 3). Two-way ANOVA with Tukey’s multiple comparison’s test used. **G** Proposed mechanism for the aberrant activation of microglia in the absence of ATM kinase. Persistent DNA damage associated with the loss of ATM, or its kinase activity, results in activation of the RELB/p52 non-canonical NF-κB pathway. This leads to microglial activation characterised by increased expression of pro-inflammatory cytokines and enhanced phagocytic clearance properties. We speculate that the release of interleukins, chemokines, and reactive oxidative species in the extracellular compartment could promote and amplify sustained non-canonical NF-κB activation via, for example, TNFR-family CD40 receptor and via NIK kinase-dependent signalling (arrows in red). Cell-intrinsic effects of ATM loss together with chronic inflammation could lead to aberrant phagocytic uptake of stressed but viable neurons and neurites. The engulfment of neuronal material could provide a secondary positive feedback loop to the non-canonical NF-κB signalling, thus contributing to progressive neuronal damage in Ataxia-telangiectasia. Positive feedback loops are shown in red arrows. Dashed arrows indicate unestablished connections. The image was made using Motifolio illustration toolkits.

To investigate the types of phagocytic events that occurred in co-cultures of WT or *ATM* KO microglia with WT or ATM-deficient neurons we used confocal microscopy. ATM-deficient microglia were found to uptake both apoptotic (positive for cleaved caspase-3) and viable (negative for cleaved caspase-3 and non-pyknotic) neuronal soma, although both types of events were relatively rare (Figure 6C). To determine whether some of the material readily phagocytosed by the *ATM* KO microglia included neurites, neuronal networks were visualised using an antibody against β3-tubulin following co-culture with CFSE-labelled microglia (Figure 6D). We observed that the *ATM* KO microglia demonstrated a significantly increased uptake of neurites (DAPI negative, β3-tubulin positive) compared to neuronal soma (DAPI positive, β3-tubulin positive) (Figure 6E). In addition, microglial clusters that formed specifically in co-cultures with neurons were detected (Supplementary Figure 7G). The number of microglial clusters that contained phagolysosomes showed a tendency to increase in co-cultures of both WT and ATM-deficient neurons with *ATM* KO, but not WT, microglia (Supplementary Figure 7H). Importantly, clusters of *ATM* KO microglia, compared with WT, were associated with disruptions in the neuronal network and damaged neurites when cultured with ATM-deficient neurons. Similar effects, albeit to a lesser extent, were observed in the case of healthy neurons (Figure 6D and 6F). Taken together, these data demonstrate the intrinsic capacity of activated *ATM* KO microglia to aberrantly engulf neuronal soma and neurites *in vitro*.

## DISCUSSION

A-T is a prototypical genomic instability disorder characterised by progressive cerebellar neurodegeneration. Neuroinflammation associated with genomic instability is increasingly recognised as a possible driver of progressive neurodegeneration (48). Previously, activation of microglia was observed in rodent models of A-T (10–12). In addition, morphological observations indicative of microglial activation were made in post-mortem brain samples from individuals with A-T (49). Here, using human cell models, we show that ATM-deficient microglia exhibit multiple features of activation, including increased expression of pro-inflammatory cytokines, cell surface CD40 and lysosomal CD68 markers and increased phagocytic clearance.

In the context of increased oxidative stress and DNA damage associated with ATM deficiency, the NF-κB proteins represent the fundamental mediators of inflammation. Surprisingly, although the canonical NF-κB pathway was inducible in both WT and *ATM* KO cells, we did not observe substantial degradation IκBα or the nuclear translocation of p65 or p50 in the absence of ATM. These data are consistent with previous observations that ATM plays multifaceted roles in activation of the IKK complex in response to genotoxic stress (30). For example, cytoplasmic ATM is essential for proteasomal degradation of IκBα via binding and recruitment of the SKP1/CULL 1/F-box protein-βTRCP (SCF^βTRCP^) E3 ubiquitin ligase complex to phosphorylated IκBα (50). In contrast, nuclear localisation of RELB, stabilisation of the NIK kinase, and proteolytic processing of p100 to p52 were apparent in ATM-deficient microglia, confirming basal activation of the non-canonical NF-κB pathway. Notably, SCF^β-TRCP^ is also involved in ubiquitin-mediated proteolysis of p100 to p52. However, the p100-SCF^β-TRCP^ complex formation was not defective in the absence of ATM (50).

At the present time, the role of non-canonical NF-κB signalling in microglia is poorly understood. We find that microglial activation in the absence of ATM, including pro-inflammatory gene expression signatures and enhanced phagocytic clearance, was dependent on RELB and mediated by NIK kinase. In ATM-proficient microglia, the RELB/p52 signalling is activated following induction of low-levels unrepaired DNA damage. The results lead us to suggest that specific activation of the RELB/p52-mediated non-canonical NF-κB pathway in microglia is linked with persistent DNA damage associated with loss of ATM function (Figure 6G). The detailed mechanistic understanding of the primary events that trigger this activation is presently unclear. One possibility is that aberrantly localised cytosolic DNA, which arises from micronuclei and damaged mitochondria in A-T (10,12,13,15), can stimulate the non-canonical NF-κB pathway. However, mechanistic links between cytosolic DNA signalling and the non-canonical NF-κB pathway are yet to be defined (48). Alternatively, poly[ADP-ribose] polymerase-1, a DNA strand break sensor, which cooperates with ATM to regulate the canonical NF-κB pathway, might play a role in RELB/p52 NF-κB signalling in A-T (51, 52).

The non-canonical NF-κB pathway promotes cell survival albeit at the expense of disease-promoting chronic inflammation and autoimmunity (35). Activation of canonical p65 signalling represses expression of anti-apoptotic genes following DNA damage (53). In addition, NIK- and RELB/p52-mediated pathways supress p65-dependent production of type I interferons and pro-inflammatory cytokines (54, 55). It is therefore possible that RELB/p52-mediated signalling in the absence of ATM provides a tolerance mechanism for limiting p65-driven pro-apoptotic responses. This mechanism is nevertheless accompanied by aberrant clearance of neurites, and to a much lesser extent neuronal soma, by *ATM* KO, but not WT, human microglia in the context of chronic inflammation. Our results indicate that highly phagocytic clusters of *ATM* KO microglia are associated with, and may drive, the damage of neuronal network via aberrant phagocytosis. Interestingly, clusters of activated microglia were previously detected in the cerebellum of A-T patients, although their function was not determined (49). Importantly, aberrant microglial pruning of neurites and synapses drives neurodegeneration in murine models of progranulin deficiency, Alzheimer’s disease and multiple sclerosis, and blocking microglial phagocytosis mitigates the pathological consequences (56–58).

The possibility that clusters of highly phagocytic *ATM* KO microglia contribute to neuronal damage via local secretion of cytokines cannot be excluded. Indeed, in murine model of A-T, microglia-induced neurotoxicity was dependent on secretion of IL-1β (12). We propose that initial activation of the non-canonical NF-κB pathway in the presence of unrepaired DNA damage could promote the release of cytokines in the extracellular compartment, thus establishing the local pro-inflammatory environment. Such inflammation could lead to cellular stress and cooperate with the aberrant phagocytic activities of microglia (Figure 6G). Our findings that activation of *ATM* KO microglia is partially rescued NIK kinase depletion confirm the contribution of cell surface receptor-cytokine interactions to inflammation. These interactions may involve the TNFR receptor CD40, expression of which is increased in *ATM* KO microglia. The result of this positive feedback loop in A-T microglia would be the amplification of non-canonical NF-κB signaling and chronic neuroinflammation contributing to disease progression. The phagocytic clearance of neuronal material would further perpetuate the non-canonical NF-κB signalling pathways and inflammation (Figure 6G).

The observation that loss of ATM drives normally neuroprotective microglia to become dysfunctional, are in general agreement with the neurotoxicity of Atm-inhibited murine microglia (12). There are, however, mechanistic differences. In mouse models of A-T, neurotoxicity is dependent on secretion of IL-1β by microglia in the absence of contact with neurons and is controlled by the STING-p65 canonical NF-κB pathway. In contrast, in human *ATM* KO microglia, we failed to observe global damage of ATM-deficient neurons via secretion. One possibility is that the long-term use of an ATM inhibitor, which acts as a dominant negative mutant of ATM (59), exacerbated the toxicity of murine microglia and contributed to neuronal susceptibility to inflammation. In the present work, the levels of phagocytosis of both damaged and live neuronal material by *ATM* KO microglia were on average 40-50% higher compared to WT cells. Although the contribution of *ATM* KO microglia to neuroinflammation and phagocytosis may appear to be modest, we suggest that even small increases in dysfunctional microglial clearance could lead to damage of PC and granule cells in a human brain. These findings are consistent with the slow progressive nature of neurodegeneration observed in individuals with A-T. Human microglia are highly immune reactive compared to their rodent counterparts (60). The differences in the threshold for execution of pro-survival strategies may explain why our findings of non-canonical NF-κB activation in ATM-deficient human microglia differ from the previously observed activation of p65 in rodent models of A-T (10, 12).

Microglia are known to dynamically interact with both the soma of PCs and their dendritic arborisations *in vivo*, and there is increasing evidence that these interactions might be dysfunctional in disease (61). For example, in a mouse model of Niemann Pick Type-C disease, which is associated with genetic defects in lysosomal storage, activated cerebellar microglia showed increased engulfment of PC dendrites, contributing to neuronal degeneration (62). Interestingly, microglia show high regional diversity, with cerebellar microglia being the most distinct based on their gene-expression signature. They exist in a highly immune-vigilant state, and this profile is further exacerbated with ageing (60). In addition, microglia in the cerebellum are enriched in genes associated with active cell clearance (63). Indeed, it is tempting to speculate that persistent DNA damage associated with the loss of ATM might exacerbate the phagocytic abilities of highly reactive cerebellar microglia *in vivo*, resulting in excessive clearance of neurites that contributes to neurodegeneration in A-T.

## MATERIALS AND METHODS

### Reagents and materials

Reagents and materials used in this work are described in Supplementary Table S1.

### Cell culture

Immortalised human fetal microglial HMC3 cells were established by Marc Tardieu and kindly provided by Brian Bigger (University of Manchester) (20). HMC3 cells were grown in DMEM (Gibco; 4.5 g/L glucose, no pyruvate) supplemented with 10% heat-inactivated fetal bovine serum (HI-FBS; Merck). Immortalized human C20 microglia were kindly provided by David Alvarez-Carbonell (21). C20 cells were grown in DMEM:F12 (Lonza) supplemented with 1% N-2 (ThermoFisher) and 1% HI-FBS. Microglia were routinely seeded at the density of 3,000-6,000 cells/cm^2^ reaching about 80% confluency at the end of an experiment.

Fetal mesencephalic LUHMES neuronal progenitors were purchased from ATCC (CRL2927) (64). Cells were grown in DMEM:F12 supplemented with 1% N-2 and 40 ng/mL recombinant zebrafish basic FGF (Hyvönen laboratory, Department of Biochemistry, University of Cambridge) in vessels coated with 50 μg/mL poly-L-ornithine (Merck) and 1 μg/mL human fibronectin (Millipore). Cells were seeded at a density of 35,000-50,000 cells/cm^2^ and differentiated for 5 days in DMEM:F12 medium supplemented with 1% N-2, 1 mM dibutyryl-cAMP (SelleckChem), 1 µg/mL tetracycline (Merck) and 2 ng/mL recombinant human GDNF (R&D Systems).

All cells were cultured at 5% CO_2_, 37°C and 95% humidity. Cell lines were routinely tested for viability and mycoplasma, and kept in culture for no longer than 40-50 population doublings.

### Establishment of knockout and rescue cell lines

*ATM* KO HMC3 cells were generated using a vector encoding SpCas9(D10A) nickase (kind gift of Feng Zhang; Addgene plasmid #48141) (65) and sgRNAs targeting *ATM* exon 4: sense_sgRNA: GATGCAGGAAATCAGTAGTT anti-sense_sgRNA: TGTGTTGAGGCTGATACATT

HMC3 cells were transfected with the vector using Lipofectamine 2000 (ThermoFisher), selected with 1 μg/mL puromycin for 5 days and single-cell cloned. *ATM* knockout C20 lines were generated using the AIO-GFP vector encoding GFP-tagged SpCas9(D10A) nickase (kind gift of Steve Jackson; Addgene plasmid #74119) (66) and sgRNAs targeting *ATM* exon 4 as above. After transfection, GFP-positive C20 microglia were sorted as single cells using the MoFlo XDP Cell Sorter (Beckman-Coulter). Successful *ATM* KO clones were identified by fragment analysis using capillary electrophoresis (Applied Biosystems) and/or immunoblotting. Gene editing was confirmed by Sanger sequencing.

Re-expression of ATM in *ATM* KO HMC3 cells was achieved using the piggyBac transgene carrying FLAG-tagged wild-type *ATM* gene. *ATM* was PCR amplified using Phusion high-Fidelity DNA polymerase (ThermoFisher) from the pcDNA3.1 Flag-His-ATM plasmid (kind gift of Michael Kastan; Addgene plasmid #31985) (67) as two individual fragments with 20-nt overhangs. The oligonucleotide sequences are shown in Supplementary Table S1 (overhangs in low case, DNA sequences complementary to the gene sequence are capitalised). The piggyBac backbone was prepared by a NotI-XhoI restriction digest using the pPB-TRE IRES-mCherry plasmid (kind gift of Bon-Kyoung Koo) (28). The fragments were assembled using NEBuilder HiFi DNA Assembly Cloning kit (New England Biolabs) to yield the pPB-TRE Flag-wtATM plasmid, and confirmed by Sanger sequencing. *ATM* KO HMC3 cells were transfected with pPB-TRE Flag-wtATM, rtTA and transposase encoding plasmids (28) at a 3:3:1 ratio using the Glial-Mag Magnetofection kit according to the manufacturer’s protocol (OzBiosciences). The cells were selected using 250 ng/mL hygromycin B for 7 days and maintained as a polyclonal population in standard growth medium supplemented with 50 ng/mL of hygromycin B. Expression of ATM was induced using 25-100 ng/mL doxycycline.

### RNA interference

Cells were transfected with 50 nM final siRNA for 72 h using Lipofectamine RNAiMax according to the manufacturer’s guidelines (Thermo Fisher Scientific). siRNA Oligonucleotides were synthesised by Merck (sequences are given in the Supplementary Table S1).

### Cell treatments

To induce NF-κB signalling, human recombinant TNFα (PeproTech) was used at 50 ng/mL for 4 h. ATM inhibitor AZD1390 (SelleckChem) was used at a final concentration of 10 nM for at least 1 h and up to 9 days (DMSO served as a control) in HMC3 cells and at 100 nM in LUHMES cells during days 2-5 of differentiation. For long-term inhibitor treatments, the medium with inhibitor was renewed every 48 h. For the inhibitor wash-out, cells were washed with PBS and cultured in the absence of inhibitor for an additional 2 days. To induce ATM-dependent phosphorylation, cells were treated with 1 µM camptothecin (CPT; Cambridge Bioscience) for 1 h. To induce apoptosis, cells were treated with 0.33 µM staurosporine (Merck) for 16 h. Treatment with 2 µM hydrogen peroxide for 5 min (H_2_O_2_; Merck) was used to induce DNA strand-breaks and oxidative damage. To induce persistent DNA damage, cells were treated with 0.5 µM etoposide, or DMSO as a control, for up to 6 days (Merck). The medium with the drug was renewed every 48 h.

### Protein extraction, biochemical fractionations and immunoblotting

Whole-cell and cytoplasmic and nuclear extracts were prepared as previously described (68, 69). Proteins were separated by 4-16% Tris-glycine SDS-PAGE, transferred onto PVDF membranes and the membranes were blocked in Intercept-TBS buffer (LI-COR Biosciences). Primary antibodies were diluted in Intercept-TBS, 0.1% Tween-20. Fluorescent secondary antibodies were used at 1:20,000 and diluted in Intercept-TBS, 0.1% Tween-20, 0.01% SDS. The antibodies are described in the Supplementary Table S1. Membranes were imaged using an Odyssey CLx system (LI-COR Biosciences).

### Quantitative RT-qPCR

RNA was extracted using an RNeasy Mini Kit as per the manufacturer’s instructions (QIAGEN). RNA integrity was verified on agarose gels by assessing the 28S:18S rRNA ratio. RNA was treated with DNAse I and reverse transcribed using qPCRBIO cDNA Synthesis Kit (PCR biosystems) according to the manufacturer’s instructions. For oligonucleotide sequences refer to the Supplementary Table S1. RT-qPCR was performed using qPCRBIO SyGreen Blue Mix Lo-ROX (PCR biosystems) and the QuantStudio 5 Real Time PCR system (ThermoFisher). The data were analysed using QuantStudio Design and Analysis software (ThermoFisher). The following cycling program was used: initial denaturation and polymerase activation at 95°C for 2 min; 40 cycles with 5 s at 95°C followed by 30 s at 65°C; followed by a melting curve step (15 s at 95°C; 1 min at 60°C; 15 s at 95°C). Ct values for all mRNAs of interest were normalised to RPS13 or IPO8 reference mRNA and the data were expressed as fold-change relative to the control using the 2^-ΔΔCt^ method.

### Comet (alkaline single cell electrophoresis) assays

Suspension cells were left untreated or treated with 2 μM hydrogen peroxide for 5 min on ice (positive control) and embedded in low-melting point agarose for 2 min on a microscope slide. The cells were lysed in buffer containing 10 mM Tris-HCl, pH 10.5, 2.5 M NaCl, 100 mM EDTA, 1% (v/v) DMSO and 1% (v/v) Triton X-100 for 1 h at 4°C. To unwind the DNA, slides were incubated in cold electrophoresis buffer containing 300 mM NaOH, pH 13, 1 mM EDTA, 1% (v/v) DMSO for 30 min in the dark and electrophoresed at 1.2 V/cm for 25 min. The samples were neutralised with 0.5 M Tris-HCl, pH 8.0 and stained with SYBR Gold nucleic acid stain (Molecular Probes) for 30 min. Samples were analysed in a blinded fashion. Percentage of tail DNA was determined for two technical replicates per sample with at least 50 cells per replicate using the CASP software (70).

### Proliferation assays

For proliferation assays, cells were labelled with 5 µM carboxyfluorescein succinimidyl ester for 8 min at (CFSE; Tonbo Biosciences) for 8 min at 37°C and seeded at progressively reduced density (no less than 3000 cells/cm^2^). Cells recovered for 24 h before samples were collected every 24 h and fixed in 70% ice-cold ethanol. CFSE fluorescence intensity was recorded for 10,000 single-cell events using a BDAccuri C6 flow cytometer (BD Biosciences). The mean of fluorescence intensity (MFI) of the green FL1-A channel (excitation 488 nm, emission 530±15 nm) was calculated using FlowJo software. Relative growth for each time point was calculated by normalising MFI^-1^ of individual time points to the MFI^-1^ value at 24 h post-seeding.

### Detection of ROS and CD40 levels by flow cytometry

To detect intracellular ROS, adherent cells were incubated with 2.5 μM H_2_DCF-DA (Molecular Probes) for 30 min. Cells were washed with PBS, collected by trypsinisation and analysed by flow cytometry on the green FL-1A channel.

For cell surface staining of CD40, live cells were incubated with anti-CD40 antibodies (GeneTex, GTX14148) for 60 min on ice, counterstained with Alexa 488 goat anti-mouse secondary antibody and analysed by flow cytometry on the green FL-1A channel. Relative intracellular ROS content and CD40 expression were calculated by normalizing MFI of each sample to the MFI value of the control.

### Transwell assays

HMC3 cells were seeded onto transwell membrane inserts with a pore diameter of 0.4 µm (Greiner) and allowed to attach for 24 h. The transwell inserts with HMC3 microglia were transferred into the wells with freshly differentiated post-mitotic LUHMES neurons that were pre-treated with 100 nM AZD1390 and cultured in 0.5% HI-FBS-DMEM or LUHMES differentiation medium supplemented with 1% HI-FBS for 48h. The number of microglia was controlled by collecting the cells from transwell inserts and counting. Neuronal health was analysed by immunofluorescence. Treatment of LUHMES cells with 0.33 µM staurosporine for 16 h served as positive control for apoptosis.

### Microglial-neuronal co-culture assays

HMC3 cells were labelled with 5 µM CFSE for 8 min at 37°C. Post-mitotic LUHMES cells were labelled with 5 µM CellTrace Violet (CTV; ThermoFisher) according to the manufacturer’s protocol. At 24 h post-labelling HMC3 microglia were trypsinised, checked for viability and seeded onto post-mitotic LUHMES neurons at a ratio of 1:5. The co-cultures were maintained in 0.5% HI-FBS-DMEM or differentiation medium supplemented with 1% HI-FBS for 24 h followed by their analyses by flow cytometry and immunofluorescence.

### Phagocytosis assays

At 24 h prior to phagocytosis assays, the growth medium on HMC3 microglia was replaced with 0.5% HI-FBS DMEM. Fluorescent carboxylated 5 µm beads (Spherotech) were added to cells at a ratio of 5:1 and incubated for 6 h at 37°C, 5% CO_2_ and 95% humidity. As a negative control, the assays were carried out in the presence of 1 µM Cytochalasin D. Cells were washed 5 times with ice-cold PBS, collected in 2% HI-FBS/PBS on ice and 10,000 single-cell events were acquired as above. The percentage of cells, which engulfed beads, was determined on the green FL-1A channel after subtracting the percentage of cytochalasin D controls (excitation 488 nm, emission 530±15 nm). Phagocytic activity was calculated by normalising MFI of all cells to MFI of a single bead.

Microglial phagocytosis of post-mitotic neurons was carried out in microglial-neuronal co-cultures as above. The cultures were washed with ice-cold 0.25 mM EDTA/PBS and collected by trypsinisation using 2% HI-FBS/PBS. Individual cell populations were gated and 10,000 events corresponding to CFSE-positive microglia were acquired on the green channel as above using an AttuneNxt flow cytometer (ThermoFisher). The percentage of phagocytic cells was calculated by identifying cells, which were positive for CTV and CFSE, on the violet channel (excitation 405, emission 450 ± 40nm).

### Immunofluorescence

Cells grown on glass coverslips were fixed in 4% methanol-free paraformaldehyde for 15 min and permeabilised in 0.2% Triton-X100/PBS for 10 min. Coverslips were blocked in 3% BSA, 0.05% Tween-20/PBS, incubated with primary antibodies for 16 h at 4°C and further incubated with fluorescently labelled secondary antibodies (1:500 dilution). Coverslips were mounted with the mounting medium (DAKO) containing 1.5 µg/mL DAPI. The antibodies are described in the Reagents and tools table.

In all immunofluorescent analyses, images of at least 5 fields of view (field of view: 318 x 318 µm) per coverslip with at least a total of 100 cells were taken in a blinded manner using an epi-fluorescent Zeiss Axio Observer Z1 (images in a single Z-plane) or a confocal Nikon Eclipse Ti (compressed Z-stack images) microscope. Confocal imaging was sequential for different fluorophore channels to obtain a series of axial images. A secondary antibody only control was used to subtract the non-specific background during quantifications. Analyses were performed using ImageJ.

To determine the levels of CD68, integrated intensity was measured using compressed Z-stack images.

The percentage of cells containing nuclear p65 or RELB staining was determined using nuclear (defined using DAPI) *vs* cytoplasmic (defined using vinculin) MFI.

Phagocytic events in microglial-neuronal co-cultures were identified using compressed Z-stack confocal images and XZ/YZ orthogonal projections. Phagosome-like structures, which were identified by the disruption of microglial CFSE staining, containing neuronal material were quantified in a randomly acquired tile of approximately 7 x 5 fields of view (2194 x 1567 µm). Events positive for both cleaved caspase-3 and CTV were considered as the uptake of apoptotic neurons, whereas events negative for cleaved caspase-3 and positive for CTV included phagocytosis of healthy cells and/or their parts. Additionally, events positive for β3-tubulin and DAPI represented the uptake of neuronal soma, whereas events positive for β3-tubulin and negative for DAPI were considered as the uptake of neurites. The relative frequency of these events was quantified. Microglial clusters were defined as a minimum of five proximal cells as above. Colocalisation of microglial clusters and the damaged neuronal network was quantified.

### Quantification and statistical analysis

Quantitative data are expressed as the mean of at least 3 independent biological experiments ± standard deviation (SD) unless stated otherwise. The data were tested for normality using the Shapiro Wilk test and the analysis informed the choice of parametric or non-parametric statistical tests. When normalisation of data to the respective control was carried out, a one-sample t-test with a theoretical mean of 1 was used. For comparison of two samples, paired or unpaired two-tailed t-test with Welch’s correction for unequal variances was used. To compare three or more unmatched samples, a one- or two-way ANOVA was used. The comparison of data obtained from different fields in immunofluorescence was carried out using a non-parametric Wilcoxon test (paired samples – variables counted from the same field). The exact *p*-values are indicated in figure legends, ns – not significant (*p* ≥ 0.05). The analyses were performed using GraphPad Prism.

## ACKNOWLEDGEMENTS

We thank Brian Ortmann and Laura Baldwin for contributing at the early stages of this work. We are grateful to Maria Daly (flow cytometry facility, MRC Laboratory of Molecular Biology, Cambridge) for technical support with single cell sorting and Richard Butler (imaging facility, Gurdon Institute, Cambridge) for support with image analysis. We thank Steve Durant (AstraZeneca) for being E.T.’s industrial co-supervisor. This work was funded by a Wellcome Trust and Royal Society Sir Henry Dale Fellowship (grant number 107643/B/15/Z), a Wellcome-Beit Prize, a Royal Society research grant (RGS\R1\201043), and an Isaac Newton Trust/Wellcome Trust ISSF/University of Cambridge Joint Research Grants Scheme. E.T. was supported by an AstraZeneca PhD studentship. Funding for open access charge: Wellcome Trust.

---

**Supplementary Figure S1:**
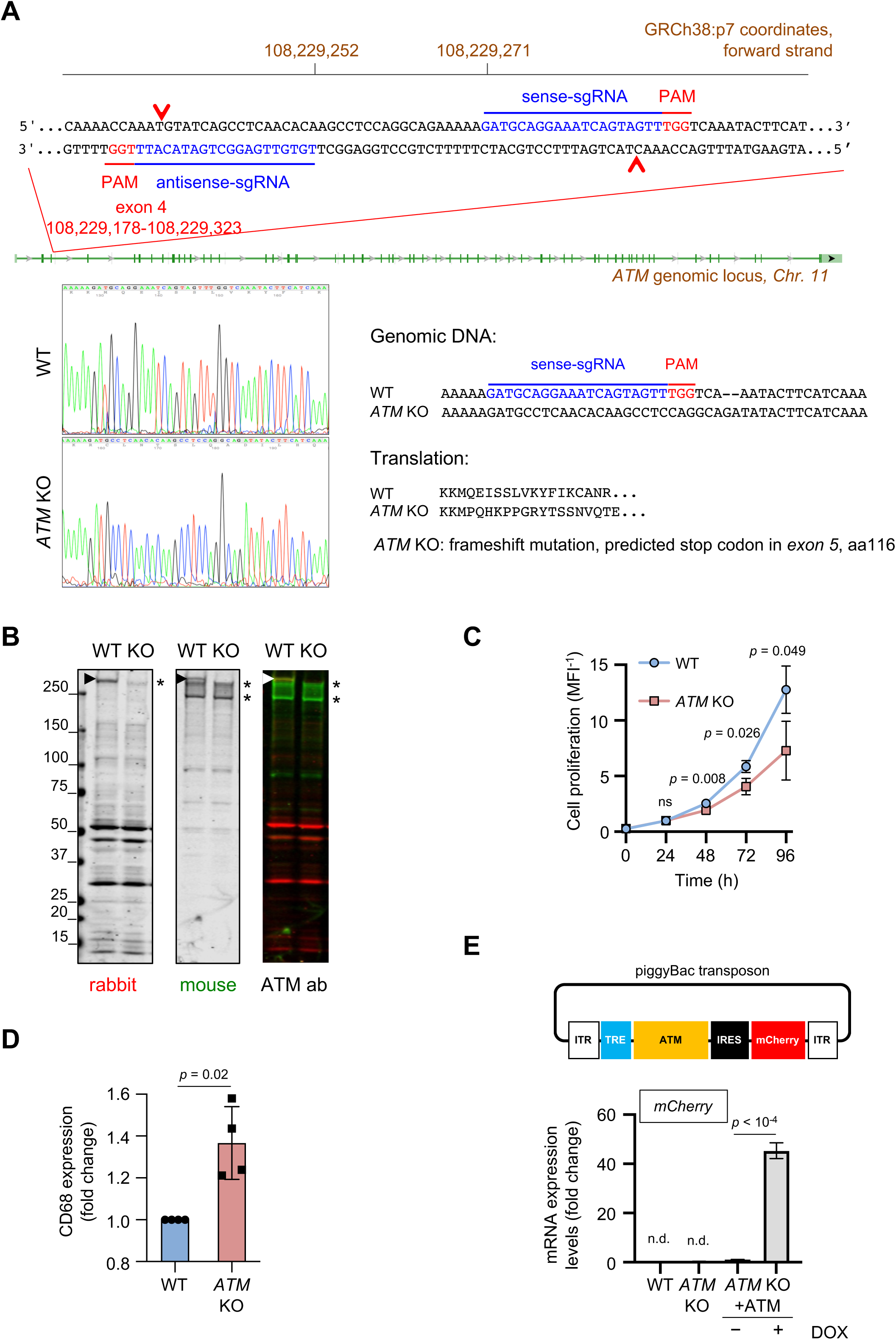
Generation and characterisation of *ATM* KO HMC3 microglia. **A** Design of CRISPR/Cas9-mediated targeting of the *ATM* locus (top). Sanger sequencing of *ATM* KO HMC3 showing 2 bp deletion in *ATM* exon 4 leading to a frame shift, premature STOP codon in exon 5 and generation of a truncated protein. WT HMC3 shown as control. **B** Immunoblot analysis of ATM protein expression in *ATM* KO vs WT HMC3. Two different ATM antibodies raised against peptides corresponding to residues 1967-1988 (mouse monoclonal, green) and 2250-2600 (rabbit polyclonal, red) in human ATM were used. **C** Proliferation curves of WT and *ATM* KO HMC3 cells. Proliferation is relative to WT HMC3 at 24 h. Mean ± S.D. shown (n = 3). Two-way ANOVA with Tukey’s multiple comparison’s test used. **D** Quantification of CD68 expression in WT and *ATM* KO HMC3 cells using immunofluorescence. Confocal Z-stack compression images were used for quantification Mean ± S.D. shown (n = 4). One-sample t-test used. **E** Schematic of piggyBac transposon carrying *ATM* gene (top) and quantitative RT-qPCR analysis of *mCherry* expression in cells as in Figure 1A (bottom). *ATM* KO cells, in which ATM was re-expressed from doxycycline-inducible piggyBac transgene, were cultured in the presence (0.1 μg/mL) or absence of doxycycline (DOX). Expression is relative to *ATM* KO HMC3, no DOX. Reference gene: *IPO8*. Mean ± S.D. shown (n = 3 technical replicates). One sample t-test used. N.d.: not detected.

**Supplementary Figure S2:**
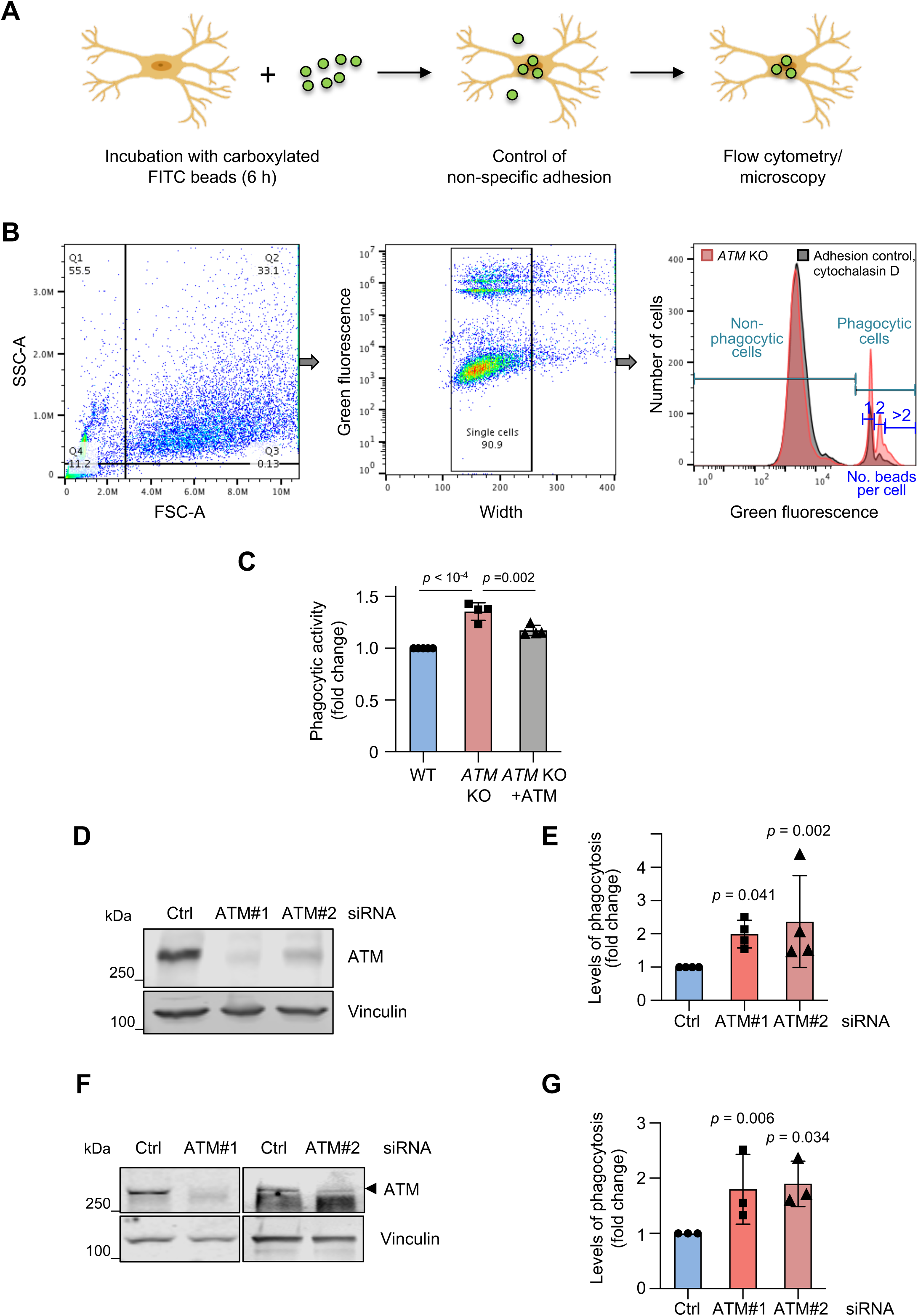
ATM-deficient microglia are activated. **A** Schematic of phagocytosis assays. **B** Workflow of phagocytosis analyses using 5 μm beads as substrates by flow cytometry. To control for non-specific adhesion, phagocytosis assays were carried out in the absence or presence of 1 µM cytochalasin D. Percentages of phagocytic cells further referred to as phagocytosis levels were determined. To calculate the relative average number of beads per cell (further referred to as phagocytic activity), mean fluorescent intensity (MFI) of phagocytic cells after subtracting MFI of the cytochalasin D control was divided by MFI of a single bead. **C** Phagocytic (5 μm beads) activity of HMC3 microglia as in Figure 1G. Activities are relative to WT HMC3. Mean ± S.D. shown (n = 4). One-way ANOVA with Tukey’s multiple comparison’s test used. **D, F** Immunoblot analysis of ATM silencing using two independent siRNA sequences (ATM#1 and ATM#2) in HMC3 and C20 microglia, respectively. Loading control: vinculin. **E, G** Phagocytosis levels (5 μm beads) of ATM-deficient HMC3 and C20 as in (**D, F**), respectively. Phagocytosis is relative to cells treated with control siRNA (Ctrl), in which 2.8 ± 2.5% (HMC3) or 11.5 ± 6.1 % (C20) of cells are phagocytic. Mean ± S.D. shown (n = 4 for HMC3, n = 3 for C20). One-sample t-test used.

**Supplementary Figure S3:**
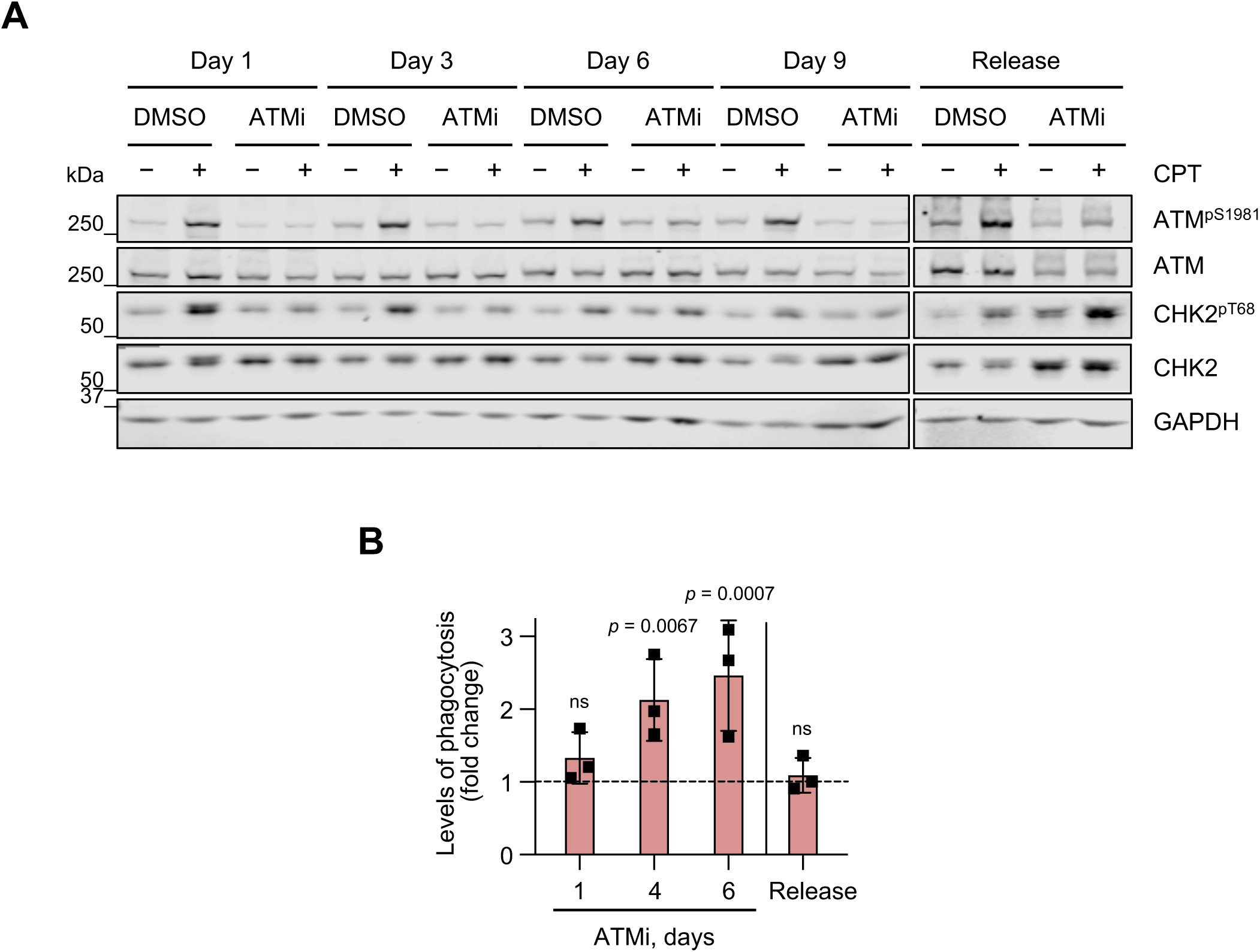
Long-term ATM inhibition leads to progressively increased phagocytosis. **A** Immunoblot analysis of the efficacy of long-term inhibition of ATM activity and inhibitor wash out in HMC3 microglia. Cells were treated with DMSO as a control or 10 nM AZD1390 for up to 9 days. The inhibitor was washed out after 6 days of inhibition for an additional 2 days (Release). To test for ATM activity, cells were treated with 1 μM camptothecin (CPT) and analysed at 1 h post-treatment. **B** Phagocytosis levels (5 μm beads) of HMC3 cells treated with ATM inhibitor as in (**A**). Phagocytosis is relative to WT HMC3 at respective time points (dashed line). Mean ± S.D. shown (n = 3). Two-way ANOVA with Tukey’s multiple comparison’s test used.

**Supplementary Figure S4:**
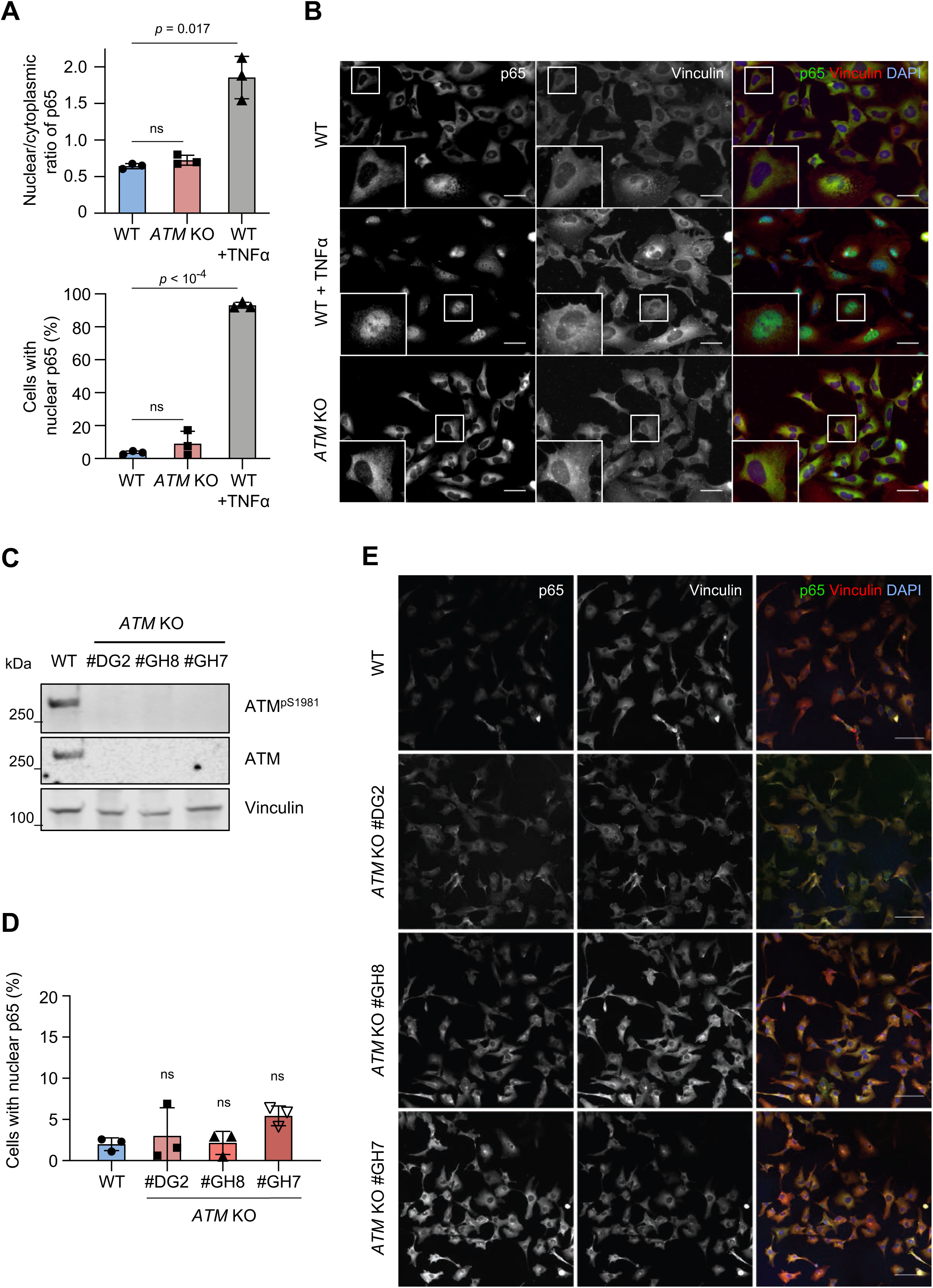
The canonical p65-dependent NF-κB pathway is not activated in *ATM* KO microglia. **A** Quantification of p65 localisation presented as nuclear/cytoplasmic ratio (top) and percentage of cells with nuclear p65 (bottom) in WT and *ATM* KO HMC3 cells. Positive control: 50 µM TNF-α for 6 h. Mean ± S.D. shown (n = 3). One-way ANOVA with Tukey’s multiple comparison’s test used. **B** Representative images of (**A**). Images in a single Z-plane shown. Scale bar: 50 µm. p65 (green), vinculin (red), DNA (DAPI, blue). **C** Immunoblot analysis of WT and three clones of *ATM* KO C20 microglia. Loading control: vinculin. **D** Quantification of p65 localisation presented as nuclear/cytoplasmic ratio of individual cells in WT and *ATM* KO C20 cells. Mean ± S.D. shown (n = 3 independent experiments). One-way ANOVA with Tukey’s multiple comparison’s test used. **E** Representative images of (**D**). Images of compressed z-stacks shown. Scale bar: 50 µm. p65 (green), vinculin (red), DNA (DAPI, blue).

**Supplementary Figure S5:**
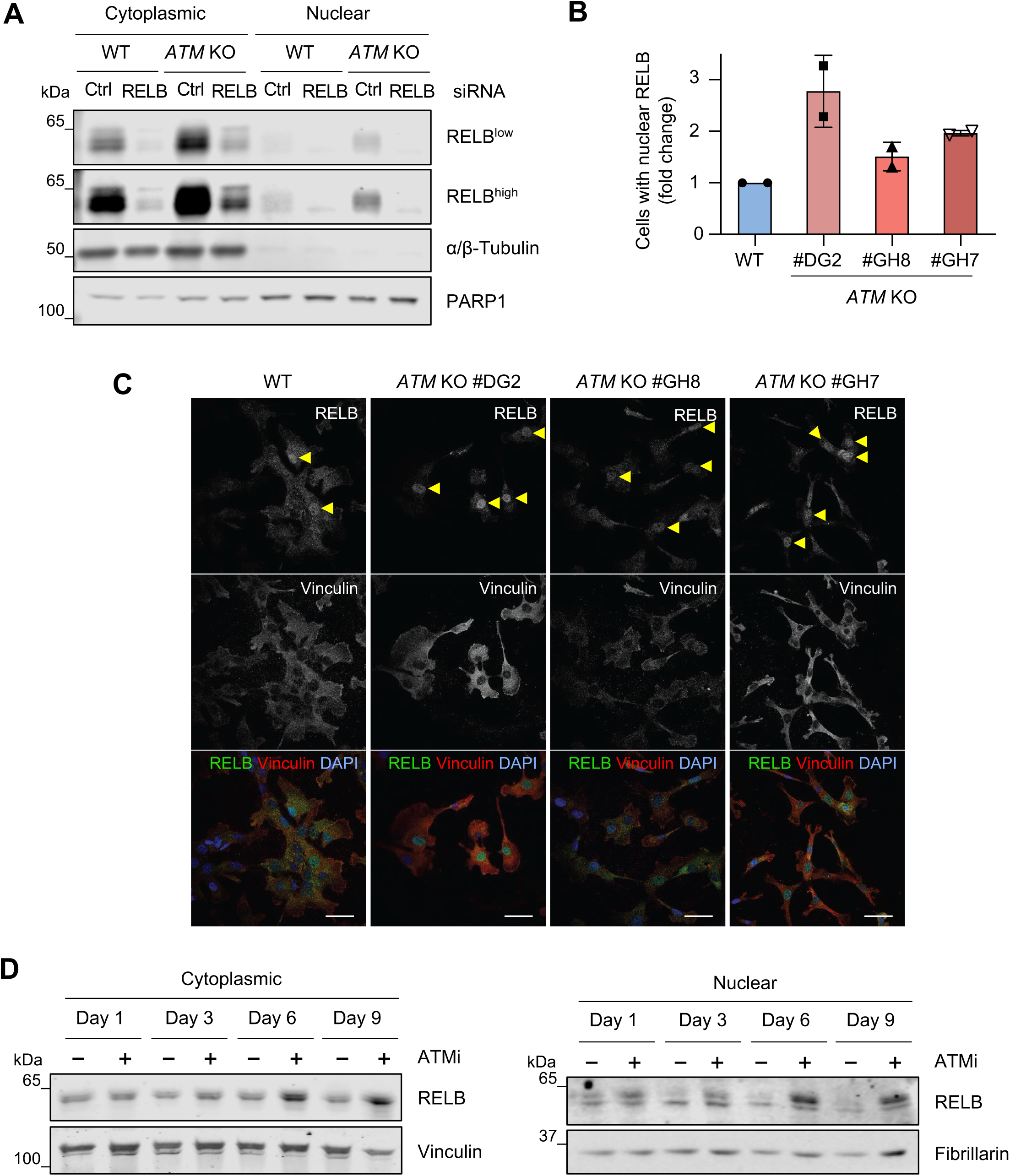
Non-canonical RELB/p52 NF-kB pathway is activated in ATM KO microglia. **A** Immunoblot analysis of siRNA-mediated silencing of RELB in cytoplasmic and nuclear extracts of WT and *ATM* KO HMC3 cells. Loading controls: a/H-tubulin (cytoplasmic), PARP1 (nuclear). **B** Quantification of RELB localisation in WT and *ATM* KO C20 cells. Data are relative to WT C20, in which 17.6 ± 12.8% of cells contain nuclear RELB. Mean ± S.D. shown (n = 2 independent experiments). **C** Representative images of (**B**). Images in a single Z-plane shown. Scale bar: 50 pm. RELB (green), vinculin (red), DNA (DAPI, blue). **D** Immunoblot analysis of RELB in cytoplasmic (left) and nuclear (right) extracts of HMC3 cells treated with 10 nM AZD1390 or DMSO control as in **Supplementary Figure S3A**. Loading controls: vinculin (cytoplasmic), fibrillarin (nuclear).

**Supplementary Figure S6:**
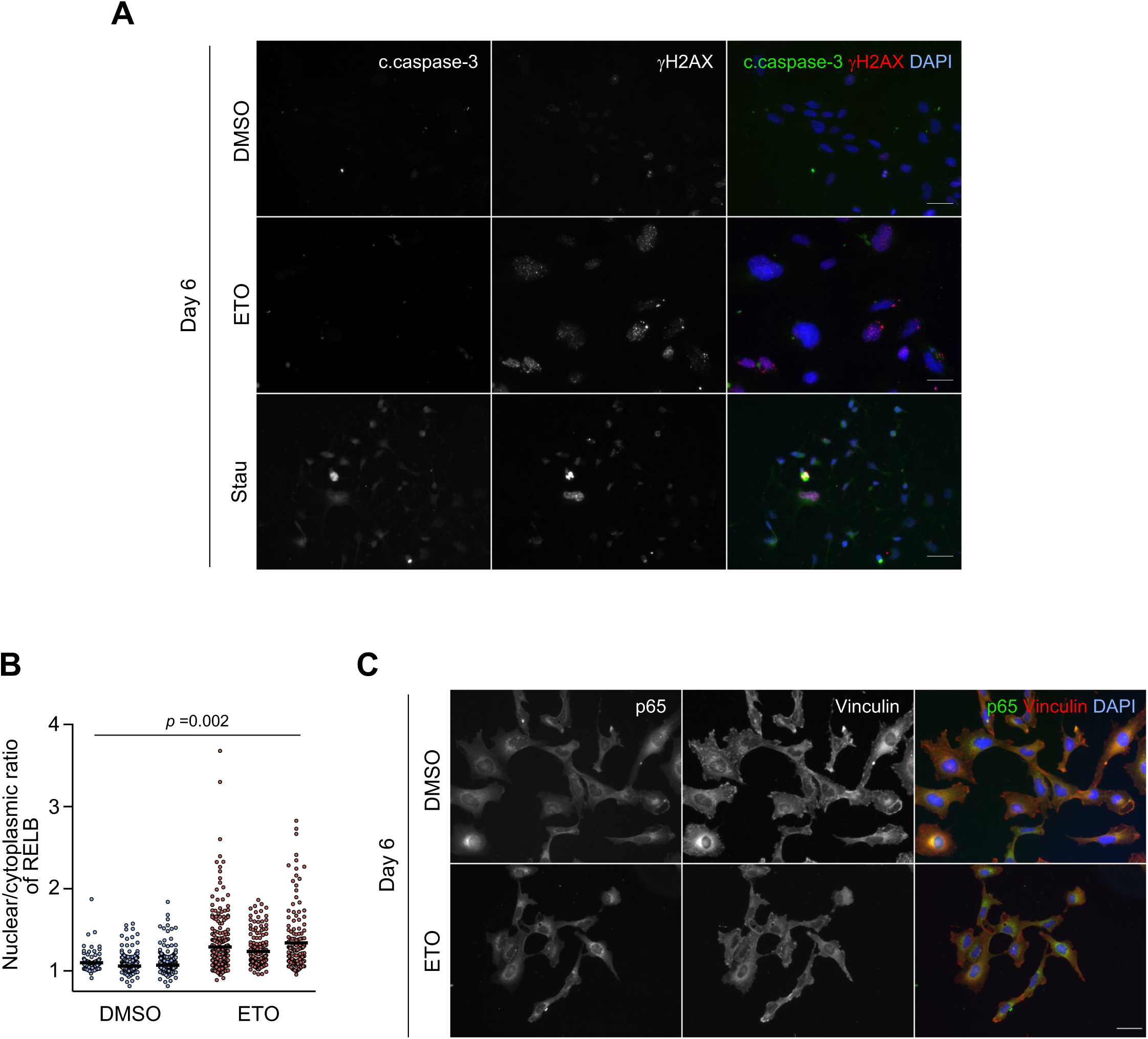
Persistent DNA damage leads to the activation of RELB/p52 NF-κB pathway. **A** Representative immunofluorescence images of HMC3 microglia treated with 0.5 μM etoposide (ETO) or DMSO as a control for 6 days as in Figure 5D, and analysed for DNA damage and apoptosis/cell death. Positive control: 0.33 μM staurosporine for 16 h (Stau). Images in a single Z-plane shown. Scale bar: 50 μm. Cleaved caspase-3 (c.caspase-3; green), γH2AX (red), DAPI (blue). **B** Quantification of RELB nuclear translocation in cells treated as in (**A**). Control: DMSO. Data presented as nuclear/cytoplasmic ratio of individual cells (100 cells per experiment; n = 3). Nested t-test used. **C** Representative immunofluorescence images of p65 localisation in cells treated as in (**A**). Images in a single Z-plane shown. Scale bar: 50 μm. p65 (green), vinculin (red), DAPI (blue).

**Supplementary Figure S7:**
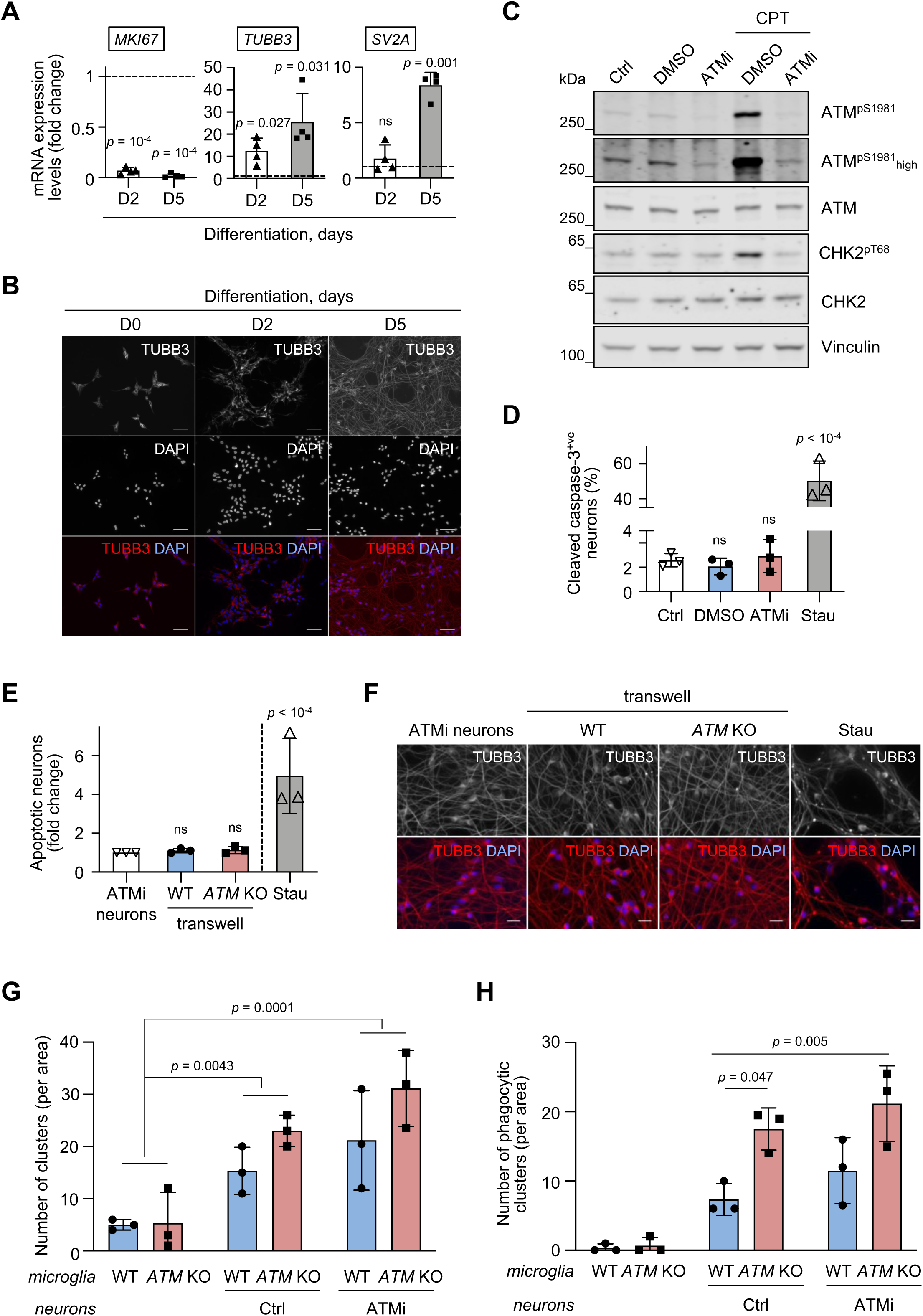
ATM deficient microglia aberrantly engulf neurites. **A** RT-qPCR analysis of proliferation MKI67, committed neuronal β3 tubulin (TUBB3) and the synaptic marker SV2A in LUHMES cells at differentiation days 2 and 5 (D2, D5). Expression is relative to day 0 (dashed line). Reference gene: GAPDH. Mean ± S.D. shown (n = 4). Unpaired t-test used. **B** Representative immunofluorescence images of cells in (**A**). Images in a single Z-plane shown. Scale bar: 50 μm. β3-tubulin (TUBB3; red), DAPI (blue). **C** Immunoblot analysis showing the efficacy and the persistence of ATM activity inhibition in post-mitotic LUHMES neurons. From day 2, cells were differentiated in the presence of 100 nM AZD1390 or DMSO for 3 days. Untreated post-mitotic neurons were used as a control (Ctrl). After 3 days of inhibition, AZD1390 was washed out and cells were cultured in standard differentiation medium for an additional 48 h. To test for ATM activity, cells were treated with 1 μM camptothecin (CPT) and analysed at 1 h post treatment. **D** Quantification of cleaved caspase-3 positive LUHMES neurons in post-mitotic LUHMES neurons treated with AZD1390 or DMSO for 3 days during differentiation. Ctrl: standard differentiation medium. Positive control: 0.33 μM staurosporine for 16 h (Stau). Data are relative to Ctrl. Mean ± S.D. shown (n = 3 for Ctrl). One-way ANOVA with Dunnett’s multiple comparison’s test used. **E** Immunofluorescence-based quantification of apoptotic (pyknotic and fragmented nuclei) post-mitotic ATM-deficient LUHMES neurons treated as in (**C**) and cultured in a transwell insert in the absence (Ctrl) or presence of WT or ATM KO HMC3 microglia for 48 h. Positive control: 0.33 μM staurosporine for 16 h (Stau). Data are relative to Ctrl, in which 9.1 ± 4.2% of cells are apoptotic. Mean ± S.D. shown (n = 3, two Stau data points are from separate experiments). One-way ANOVA with Dunnett’s multiple comparison’s test used. **F** Representative immunofluorescence images of LUHMES cells in (**E**). Images in a single Z-plane shown. Scale bar: 20 μm. β3-tubulin (TUBB3; red), DAPI (blue). **G** Quantification of clusters formed by WT or *ATM* KO HMC3 microglia on their own or in co-culture with post-mitotic LUHMES neurons that were pretreated with AZD1390 as in (**C**) or left untreated (Ctrl). Mean ± S.D. shown (n = 3). Two-way ANOVA with Sidak’s multiple comparison’s test used. **H** Quantification of phagocytic microglial clusters from (**E**). Mean ± S.D. shown (n = 3). Two-way ANOVA with Tukey’s multiple comparison’s test used.

**Supplementary Table S1.**
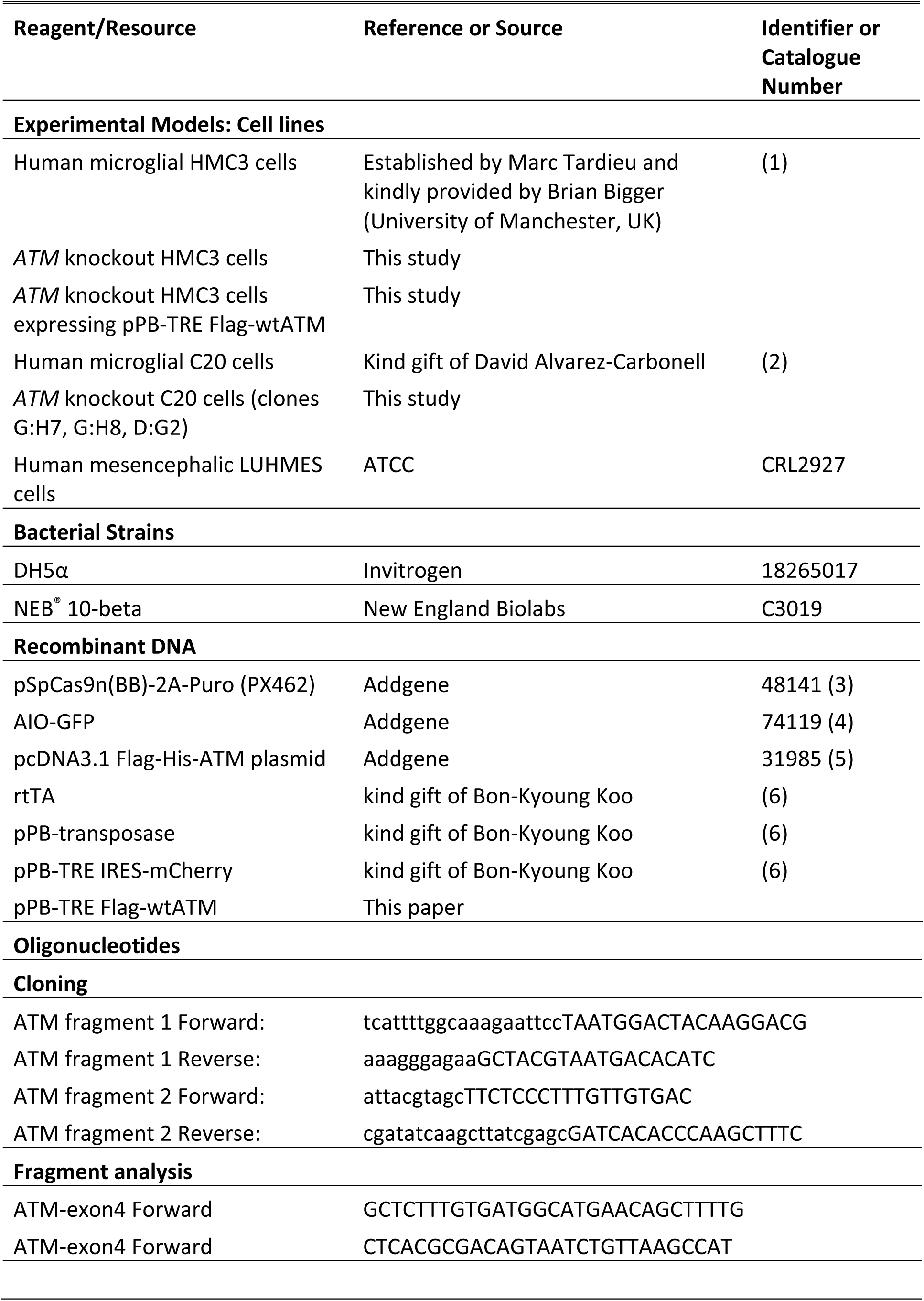
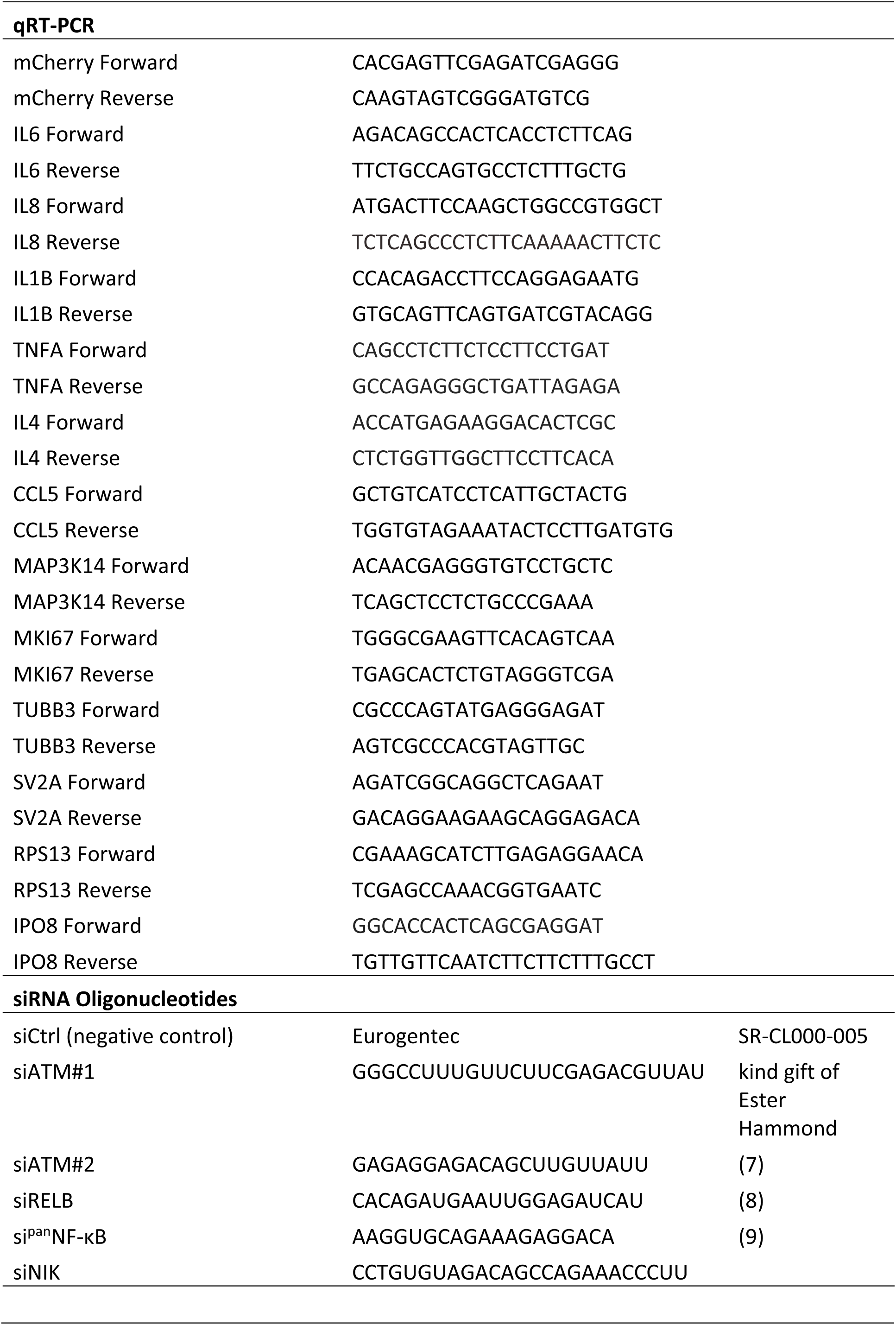
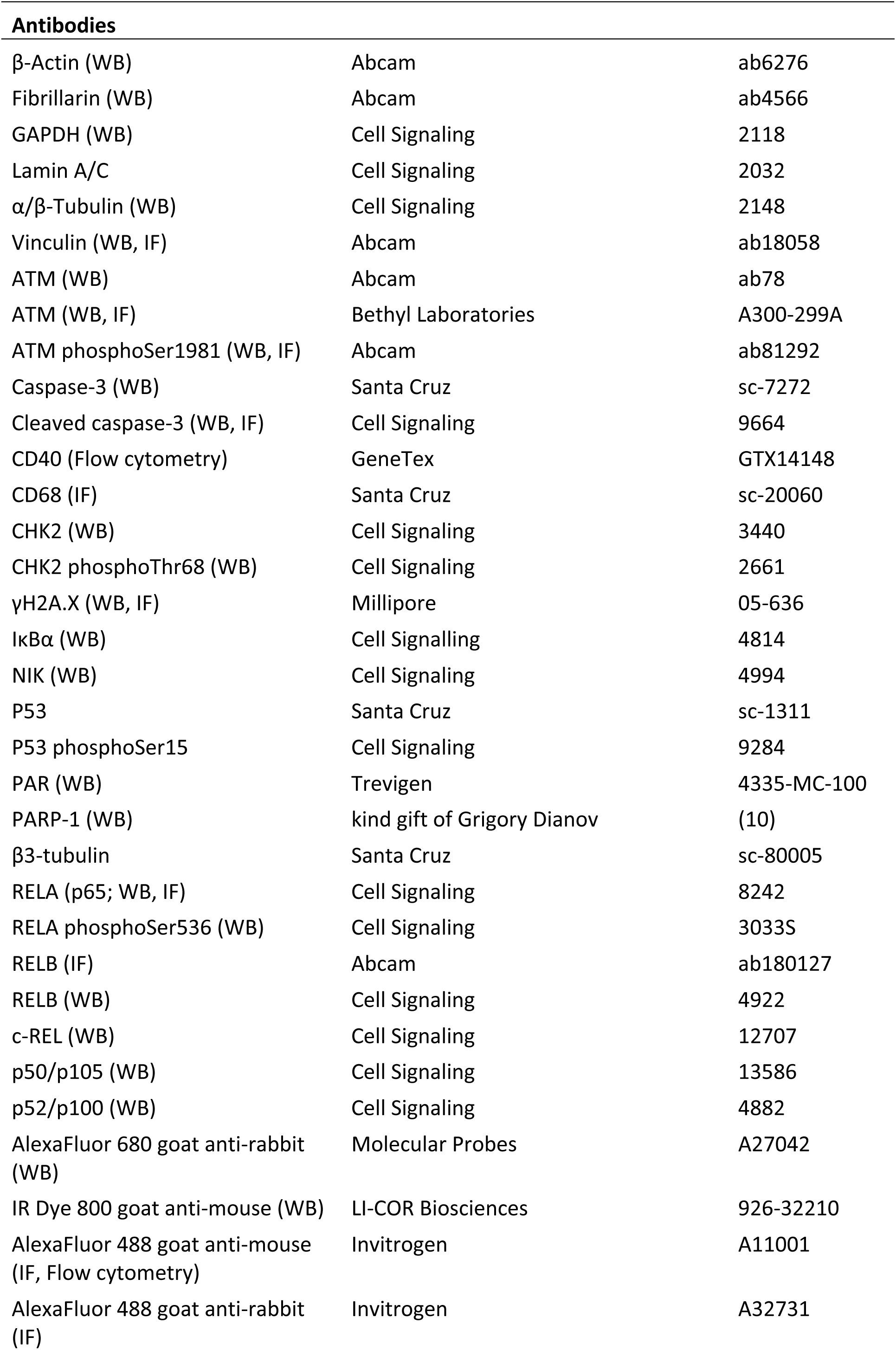
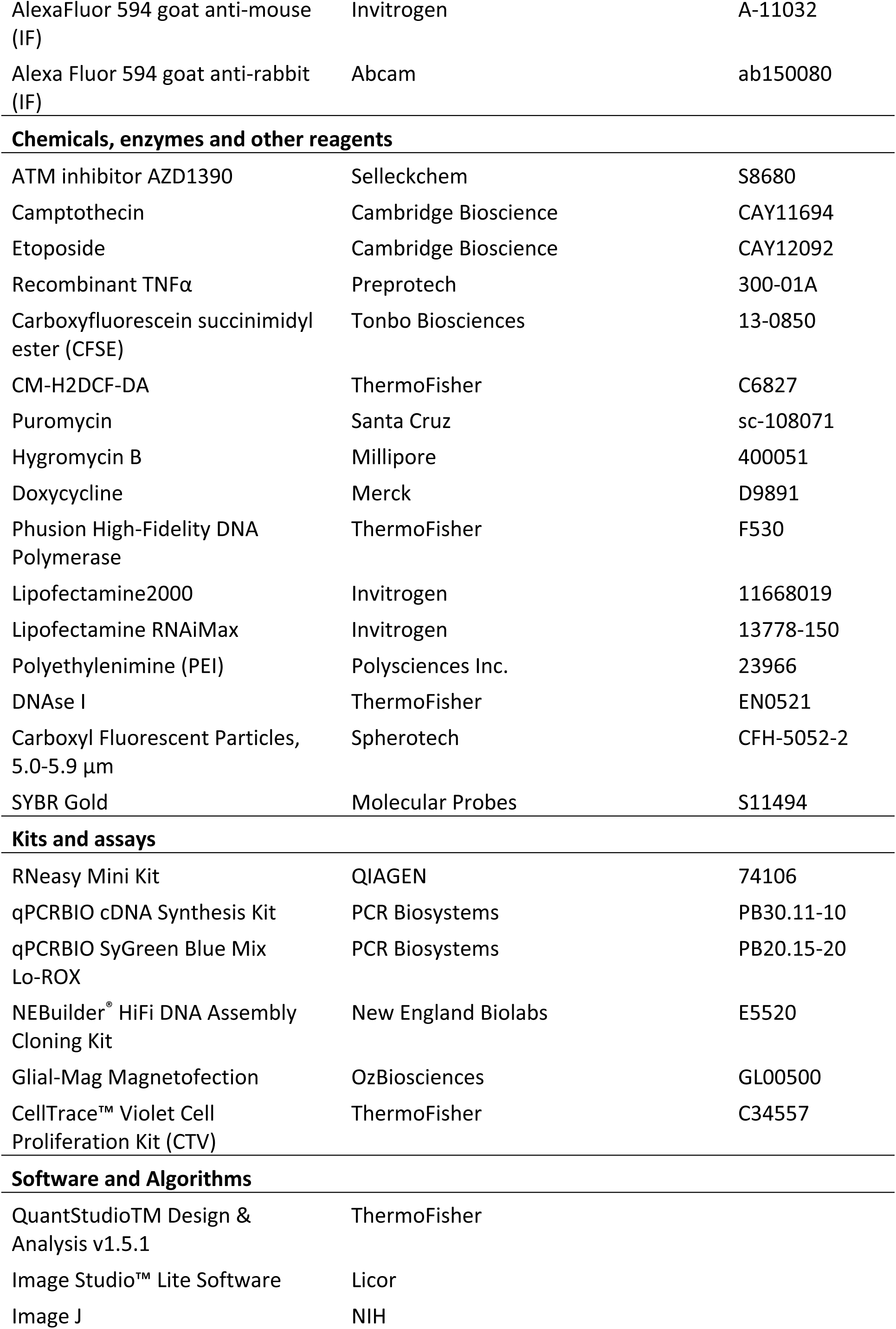
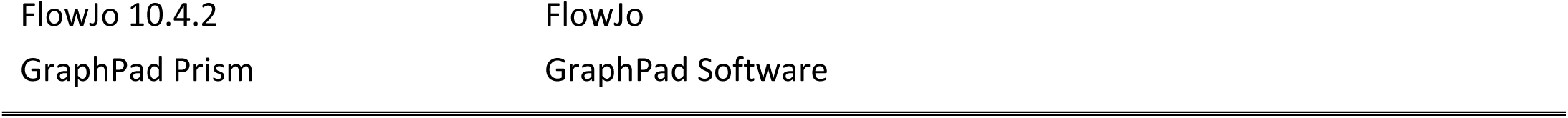
Reagents and tools used in the study.

